# High-order mutants reveal an essential requirement for peroxidases but not laccases in Casparian strip lignification

**DOI:** 10.1101/2020.06.17.154617

**Authors:** Nelson Rojas-Murcia, Kian Hématy, Yuree Lee, Aurélia Emonet, Robertas Ursache, Satoshi Fujita, Damien De Bellis, Niko Geldner

**Affiliations:** Department of Plant Molecular Biology, Biophore, Campus UNIL-Sorge, University of Lausanne, CH-1015 Lausanne, Switzerland; Umeå Plant Science Centre, Department of Plant Physiology, Umeå University, SE-901 87 Umeå, Sweden; School of Biological Sciences, Seoul National University, Seoul 08826, Republic of Korea; Plant Genomics and Breeding Institute, Seoul National University, Seoul 08826, Republic of Korea; National Institute of Genetics, 1111 Yata, Mishima, Shizuoka 411-8540, Japan; Electron Microscopy Facility, University of Lausanne, 1015 Lausanne, Switzerland

## Abstract

The invention of lignin has been at the heart of plants’ capacity to colonize land, allowing them to grow tall, transport water within their bodies and protect themselves against various stresses. Consequently, this polyphenolic polymer, that impregnates the cellulosic plant cell walls, now represents the second most abundant polymer on Earth, after cellulose itself. Yet, despite its great physiological, ecological and economical importance, our knowledge of lignin biosynthesis *in vivo*, especially the crucial last steps of polymerization within the cell wall, remains vague. Specifically, the respective roles and importance of the two main polymerizing enzymes classes, laccases and peroxidases have remained obscure. One reason for this lies in the very high numbers of laccases and peroxidases encoded by 17 and 73 homologous genes, respectively, in the Arabidopsis genome. Here, we have focused on a specific lignin structure, the ring-like Casparian strips (CS) within the endodermis of Arabidopsis roots. By reducing the number of possible candidate genes using cellular resolution expression and localization data and by boosting the levels of mutants that can be stacked using CRISPR/Cas9, we were able to knock-out more than half of all laccases in the Arabidopsis genome in a nonuple mutant – abolishing the vast majority of laccases with detectable endodermal-expression. Yet, we were unable to detect even slight defects in CS formation. By contrast, we were able to induce a complete absence of CS formation in a quintuple peroxidase mutant. Our findings are in stark contrast to the strong requirement of xylem vessels for laccase action and indicate that lignin in different cell types can be polymerized in very distinct ways. We speculate that cells lignify differently depending on whether they deposit lignin in a localized or ubiquitous fashion, whether they stay alive during and after lignification as well as the composition of the cell wall.

## INTRODUCTION

Casparian strips are highly conserved structures that are a defining feature of the root endodermis in higher plants. That CS are of a lignin-like nature had been proposed repeatedly since their discovery in the 19^th^ century and was firmly established by modern histological and genetic analyses in the model plant Arabidopsis (1,2). CS are strictly localized cell wall impregnations, forming as longitudinal, centrally located belts between endodermal cells. Their highly coordinated and simultaneous appearance in endodermal neighbors leads to the fusion of these belts into a supracellular structure that takes the appearance of a delicate network, due to the very thin primary cell walls of young endodermal cell (100-200 nm in width). Using lignin stains, this fine network can be easily overlooked, next to the much more pronounced lignification occurring at the same time in the thick secondary cell walls of protoxylem vessels, only two cell layers below within the vasculature. The CS therefore represent only a minor fraction of the overall lignin content of a root and are always found in close association to the xylem, making it very difficult to use the classical chemical methods of lignin analysis that are central to the field. Yet, CS are very attractive for cell biological and genetic analyses for a number of reasons. The endodermis can be observed in very young, 5-day-old seedlings and it is a relatively peripheral, large cell type, that is more easily observed than cell types in the vasculature. Moreover, its highly predictable and restricted lignification allows for meaningful spatial correlations between protein localization and lignin deposition. Finally, the endodermis stays alive during and after the entire process of lignification (3). Using the endodermis as a model, we were able to establish a strong requirement for reactive oxygen species (ROS) production in CS lignification. Knock-outs of a single NADPH oxidase, RBOHF, led to a near absence of lignification in the endodermis, as did inhibitor treatments interfering with ROS production or accumulation (2). Intriguingly, RBOHF specifically accumulates at the site of CS formation, which is initiated by the accumulation of CASPARIAN STRIP MEMBRANE DOMAIN PROTEINS (CASPs), small transmembrane scaffold proteins that are thought to recruit RBOHF and other proteins to their site of action. Another class of proteins that appear to be localized by CASPs are type III peroxidases. Among those, PER64 showed an especially strict co-localization with CASP1. Indeed, T-DNA and amiRNA driven knock-out/knock-down of multiple endodermis-expressed and CS-localized peroxidases led to a delay of barrier formation (2), although the severity and nature of lignin defects where not assessed at that time. These findings led to a model whereby CASPs are acting to bring together NADPH oxidase and peroxidase, effectively allowing to channel localized ROS production toward peroxidases, thus ensuring localized and efficient lignification (2). Clearly, localized presence of RBOHF and peroxidases is sufficient for localizing lignification, since complementation of monolignol-deficient plants with large amounts of external monolignols, did not affect localization of CS formation (1). More recently, we showed that a dedicated receptor pathway in the endodermis can detect defects in the CS diffusion barrier and initiate compensatory, ectopic lignification in cell corners, both by enhancing ROS production from RBOHF and RBOHD and inducing expression of genes, including additional peroxidases (*PER*) and laccases (*LAC*) (4–6). Our findings demonstrating a strong requirement for ROS production and a partial genetic requirement for peroxidases was in contrast with the finding that LACCASES are necessary for lignification of xylem vessels, fibers and other cell types. LACCASEs (LAC) are glycosylated, multi-copper enzymes that catalyze the oxidation of various phenolic substrates using O_2_ as final electron acceptor, not requiring H_2_O_2_ (7,8). LAC15 was found to express in seed coats, its loss-of-function mutant displaying a 30% reduction of lignin in seeds (9,10). *LAC4*, also called *IRREGULAR XYLEM 12* (11) and *LAC17* are expressed in vessels and xylary fibers in the stem (12). LAC4 localizes to secondary cell wall domains of proto- and meta-xylem vessels, as well as vessels, xylary and interfascicular fibers of inflorescence stems (12–15). Mutant lines of *lac4* and *lac4 lac17* have collapsed xylem vessels and reduced lignin content in total stem biomass (12). The reduction in lignin content of the double mutant was further enhanced in the presence of a *lac11* mutant, which showed severe developmental defects and stopped growing after developing the two first pair of leaves (16). Finally, *LAC15* and *LAC7* are expressed in lignifying cells in the context of floral organ abscission, although it was not demonstrated whether mutations of both *LACs* affected lignin deposition in this context (17). More recently, *LAC2* was found to negatively regulate overlignification in the root vasculature upon phosphate and water deficiency. Nevertheless, the exact role of LAC2 in such context was not determined (18).

Yet, numerous studies have also provided evidence for participation of PERs in lignification outside of the endodermis. In the *Arabidopsis* stem xylem vessels, mutation of *PER*s partially impact lignification (19,20). It was proposed that both enzymes act sequentially in the lignin polymerization of a same cell type, although it has not been excluded that they individually lignify different cell types (8,15,21). Cell cultures of the gymnosperm Norway spruce release lignin polymers into the growth medium (22). Scavenging of H_2_O_2_ prevents production of this extracellular lignin and phenolic profiling of the culture media of scavenger-treated cell cultures revealed the accumulation of specific oligolignols (21). This led the authors to propose that H_2_O_2_-independent enzymes, such as laccases, mediate formation of oligolignols, while peroxidases would be required for further polymerization. However, the authors could not exclude the possibility that in their treatment to scavenge H_2_O_2_ a residual PERs activity was present (21). Unfortunately, participation of PER in lignification *in planta* has often been inferred from the use of inhibitors of PERs or H_2_0_2_ production (2,23), while genetic evidence could reveal only weak, partial effects on lignification, allowing for the possibility that peroxidases are only peripheral actors of lignification *in planta*.

In addition to PERs, data of cell type-specific gene expression has revealed that *LACs* are expressed in the endodermis (24–27). Considering the evidence on the role of *LACs* in lignification, we took advantage of the experimental setup offered by the endodermis to conduct a genetic analysis of *LACs* in order to determine whether they contribute to lignin polymerization in the CS.

In this work, we set out to determine whether one given lignin structure in a cell requires either laccases or peroxidases exclusively, or whether its formation requires a combination of both enzymes classes. Using the endodermal CS as a model, we demonstrate that a number of laccases show specific expression in the endodermis and localization to the CS, yet generation of a nonuple mutant, knocking-out the vast majority of laccases with detectable endodermal expression, had no discernable effect on CS lignification or formation of the endodermal barrier. By contrast, generating a quintuple knock-out of endodermis-enriched peroxidases led to a complete absence of CS lignification. Abrogating the compensatory lignification by the SCHENGEN pathway even led to a complete absence of any lignification in young endodermal cells.

Based on this, it is most parsimonious to conclude that, despite their strong presence, laccases are fully replaceable for lignification of CS, while peroxidases are absolutely required.

## RESULTS

### Selection of candidate *LACs* for endodermal Casparian strip formation

A reverse genetic analysis of *LACs* involved in lignification of the CS is challenging due to the fact that there are 17 encoding *LAC* genes in *Arabidopsis* with evidence for functional redundancy between them (12,16). Data of cell-type specific gene expression available for the *Arabidopsis* root is a powerful tool to guide such an effort (26–28). We found that a set composed of *LAC1, 3, 5, 13* and *16* displayed enriched mRNA expression in the endodermis (Fig. S1A-B). *LAC8* displayed strong expression across different cell types/root zones, including the endodermis (Fig. S1A-B), but since it was not enriched in endodermis, we decided to use *LAC1, 3, 5, 13* and *16* in the first step.

### *LAC1, LAC3, LAC5* and *LAC13* promoter reporters corroborate endodermal expression

We generated transcriptional reporters of *LAC1, 3, 5, 13* and *16* by fusing their upstream regulatory sequences to an NLS-GFP-GUS reporter gene. We observed expression in the endodermis for all lines except *pLAC16*, which could not be detected either as GFP, or after extended GUS staining. CS formation/lignification occurs in a reproducible spatial pattern, starting between 6-8 cells after onset of elongation (exit from the meristem). After rapid progression into an uninterrupted network, CS slowly broaden and lignin stains increase in intensity until a region of about two third of the root length - in 5-day old seedlings - at which CASP proteins start to disappear and suberin lamellae start to sever the direct contact between the CS and the plasma membrane. At this point, it can be presumed that CS lignification has ceased completely. Genes involved in CS lignification can therefore be predicted to follow this spatial pattern, an aspect that can be assessed by promoter lines, but is insufficiently resolved in cell-type-specific expression databases.

We found that expression in the *pLAC1* line was observed 6 cells after onset of elongation, starting in the stele. Only at about 16 cells, a GFP signal could additionally be observed in the endodermis (Fig. 1A and S2A). In later root developmental stages, expression became exclusive to the endodermis extending until two thirds of the total primary root length.

**Figure 1.**
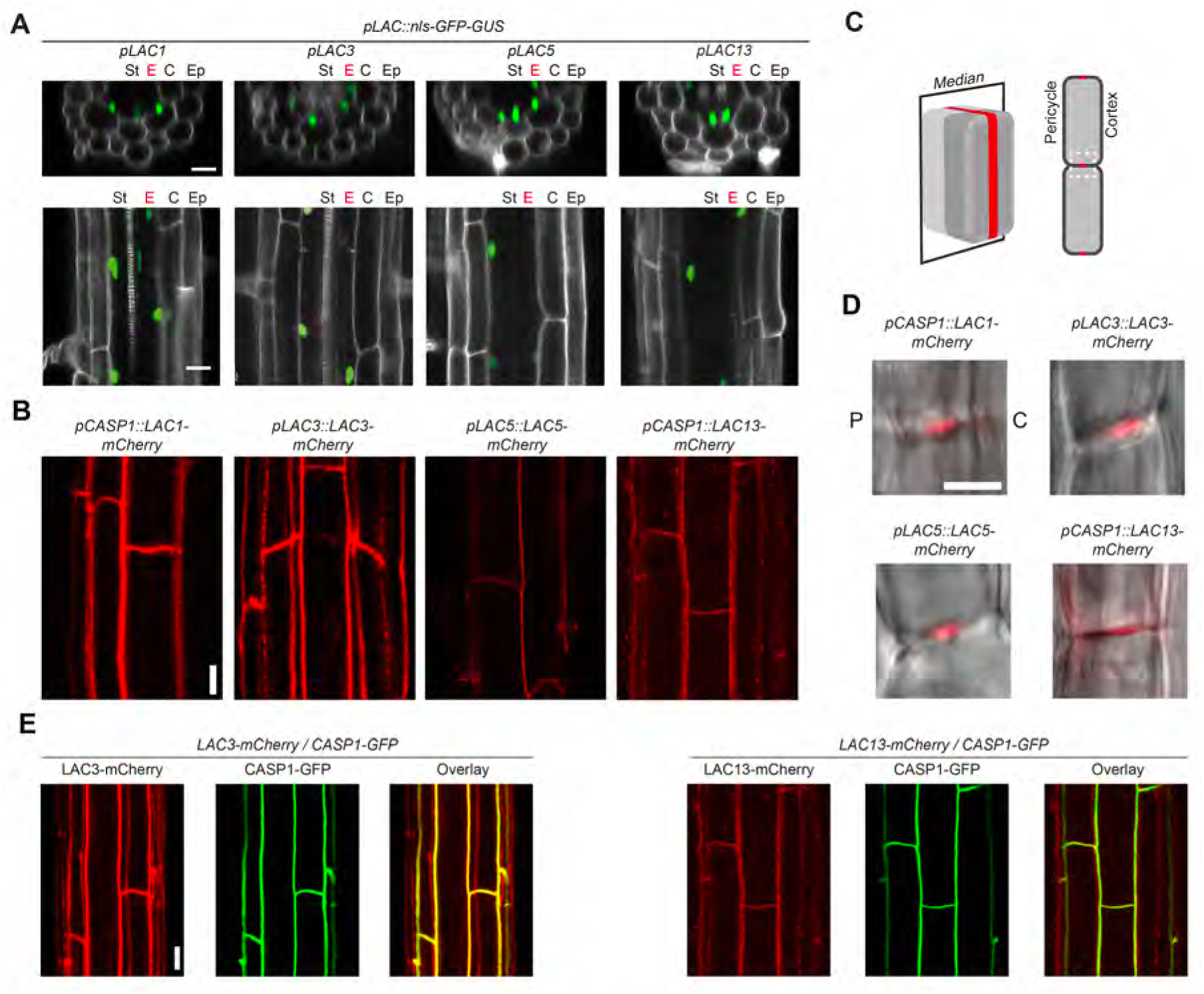
LACs are expressed in the endodermis and targeted to the Casparian strip. **(A)** Overview of expression pattern of *pLACs::nls-GFP-GUS* transcriptional reporters in 5-day-old roots of *A. thaliana* for *LAC1, LAC3, LAC5* and *LAC13.* Pictures show the overlay of GFP (green) and propidium iodide (gray). Top panel, optical transversal sections. Bottom panel, optical longitudinal sections. St, stele. E, endodermis (in red to emphasize its position). C, cortex. Ep, epidermis. **(B)** Surface overview (z-axis max. projection) of C-terminal, mCherry-4G tagged proteins of *LAC1, LAC3, LAC5* and *LAC13* localization in the endodermis. **(C)** Schematic of endodermal cell, explaining optical section shown in **D.** Schematic adapted from (40). **(D)** The 4 LACs localize to the median position of the endodermis cell wall where the CS is located. Overlay of mCherry-4G signal in red with transmitted light in grey, highlighting the cell contour. C, Cortex. P, pericycle. Other legends as in A. **(E)** C-terminal, fluorescently tagged proteins *pLAC3::LAC3-mCherry-4G* and *pCASP1::LAC13-mCherry-4G* colocalize with CASP1-GFP, which marks the membrane domain of the CS. Scale bars 20 μm in **A**; 10 μm in **B**; 5 μm in **D**; and 10 μmin **E**.

GFP expression in *pLAC3, pLAC5* and *pLAC13* lines was observed exclusively in the endodermis (Fig. 1A). However, their onset of expression varied among the three promoters, taking place after ca. 6, 9 and 25 endodermal cells for *pLAC3, pLAC5* and *pLAC13*, respectively. Additionally, while *pLAC3* and *pLAC13* expression was strong all along their expression domain, *pLAC5* expression became stronger in later root developmental stages (Fig. S2A). Expression in all three promoter lines extended until the hypocotyl.

A previous report using GUS transcriptional reporters showed that a *pLAC*7 line drove strong expression in the root (29). We generated a *pLAC7::nls-GFP-GUS* line and found strong expression of GFP in epidermis and cortex, but not in the endodermis (Fig. S2B). Taken together, the identified *LAC* promoter activities nicely matched the developmental zones where lignin deposition is observed in the CS (30) confirming and extending on the available public expression profile databases.

### Protein fusions of LAC1, LAC3, LAC5 and LAC13 accumulate at the site of Casparian strip formation

The strict, CASP-dependent localization of lignification within endodermal cell walls allows to predict that enzymes involved in lignin polymerization should accumulate at CS and co-localize with CASPs. We therefore generated plants expressing C-terminal, mCherry fusion constructs, under the control of their own, or of the strong, endodermis-specific CASP1 promoter (*pCASP1*) (31) and acquired longitudinal median and surface sections of endodermal cells on 5-day-old seedlings.

LAC1-mCherry fusion protein expressed under the control of the CASP1 promoter was observed in endodermal cells (Fig. 1B) where it localized to the median position of the endodermal cell walls as well as to the edge of the cortex-facing cell wall (Fig. 1D and S3B).

LAC3-mCherry protein could also be localized using its endogenous promoter and localized in a manner very similar to LAC1-mCherry (Fig. 1B). LAC5-mCherry-4G localized to the CS position in the cell wall and to the edges of cortex-facing cell wall of the endodermis (Fig. 1D and S3B).

LAC5- and LAC13-mCherry also accumulated preferentially at CS. Accumulation in cortex-facing cell wall corner and, in the case of LAC13, intracellular, mobile compartments, might be due to overexpression by the strong CASP1 promoter (Fig. 1B,D and S3B).

Finally, we attempted to localize LAC16-mCherry by expressing it from the strong *ELTP* promoter, active in differentiated endodermis (32). However, we could only observe a fuzzy, cytoplasm-like signal. This would indicate that the LAC16 fusion protein is not secreted. We propose that the very weakly expressed LAC16 might be a non-functional pseudogene (Fig. S2C).

To ascertain that laccases indeed localize precisely to incipient CS, we co-localized LAC3- and LAC13-mCherry with CASP1-GFP, which localizes to the plasma membrane subjacent to the CS, called the CS membrane domain (CSD). Both proteins showed near-perfect co-localization at the median position (Fig. 1E). CASPs co-localize with other cell wall proteins at the CS, such as PEROXIDASE 64 (PER64) and ENHANCED SUBERIN 1 (ESB1), both involved in CS formation (2,33). Thus, laccases and peroxidases localize to the same cell wall domain in the endodermis. This contrasts with a previous report, in which fluorescently tagged LAC4 and PER64 were localized to distinct cell wall domains in interfascicular fibers, with LAC4 localizing to the thick secondary cell wall, whereas PER64 localized to the cell wall corners and middle lamellae (15). In the endodermis, by contrast, where there is no apparent secondary cell wall, PER64 and LAC3 appear to reside in the same cell wall domain. Taken together, we found that LAC1, LAC3, LAC5 and LAC13 localize to the central median region of the endodermal cell wall, where the CS is formed.

### *A LAC1, 3, 5*, 7, 8, 9, 12, *13, 16* nonuple mutant does not have any discernable defect in Casparian strip formation

In order to test whether the LACs with enriched endodermal expression and CS localization are indeed required for lignin polymerization in the CS, we generated a *lac1 lac3 lac5 lac13 lac16* quintuple mutant (hereafter referred to as *5x lac*), by combining T-DNA insertion alleles for *LAC1, LAC5, LAC13* and *LAC16* (*lac1, lac5, lac13 lac16* hereafter referred to as *4x lac*) into which we introduced a *lac3* knock-out by CRISPR-Cas9 mutagenesis (34,35), targeting the first exon of *LAC3* (Fig. S4). Single *lac3* alleles were also generated and tested (Fig. S3C). To our surprise, we were unable to observe any difference in signal strength, structure or position of the CS, as visualized by histological stains (Figure S5A).

Since both, the specific expression pattern, as well as their specific subcellular localization, strongly suggested an important function of laccases in CS formation, we decided that the most probable explanation for an absence of defects is a higher-than-suspected degree of redundancy among the laccases. We therefore selected an additional set of four *LACCASE* genes, based on published, cell-type-specific RNA profiling datasets, focused on the endodermis (27,28). Moreover, we integrated recent RNA profiling data after SCHENGEN pathway stimulation in which LACCASE 12 was found to be upregulated, possibly compensating for a lack of activity in the *5x lac* mutant. Using a multiplex guideRNA cloning strategy we used 6 sgRNA targeting 4 LAC genes. We used two guides for LAC7, two for LAC8 (one targeting also LAC9) and two guides for LAC12 (Fig. S4B). With this multi-sgRNA construct, we were able to obtain knock-out alleles of LAC 7, 8, 9, 12 in the *5x lac* background, generating a *9x lac* mutant (Fig. S4C). Yet, despite having knocked-out the majority of *LACCASE* genes in the Arabidopsis genome – and all *LACCASES* with a significant, detectable expression in the endodermis – we were again unable to observe even a slight, quantitative difference in onset, strength, position or function of CS in the endodermis (Fig. 2). Even when we challenged the endodermis with CIF2 peptide (a ligand of the SGN3 receptor that induces overlignification in cell corners (6), we could not observe any delay, or weaker accumulation of lignin (Fig. S5B,C), demonstrating that LACCASES are also not required for the less organized, cell corner lignification in the endodermis.

**Figure 2.**
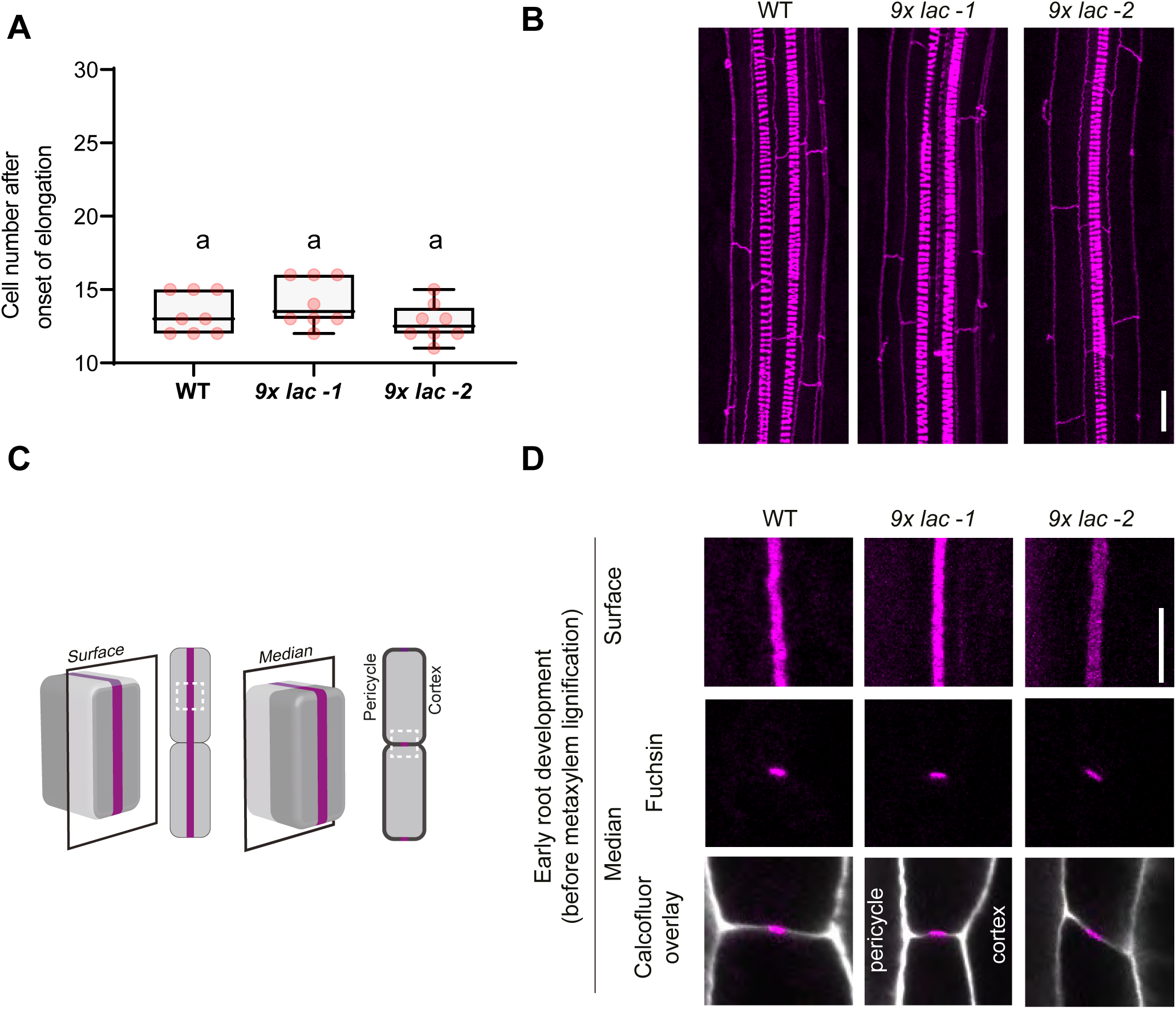
Nonuple laccase mutants *lac1;3;5;7;8;9;12;13;16* (*9x lac*) deposit functional and lignified CS. **(A)** The CS in *9x lac* (nonuple) mutants still effectively seals the apoplast. Apoplastic barrier function of the endodermis is tested by the propidium iodide diffusion assay. Numbers indicate the endodermal cell in which apoplastic block is established, counting from the onset of endodermis cell elongation (one-way ANOVA and Tukey’s test as post hoc analysis show no statistically significant difference). WT, wild-type Col-0. **(B)** Basic fuchsin staining for visualization of lignin deposition in roots of two *9x-lac* mutants compared to WT. Pictures show confocal z-axis maximum projections along the root, visualizing the CS networks and xylem vessels. Note the continuous, uninterrupted CS of the nonuple mutants. **(C)** Schematic of endodermal cell, explaining optical sections shown in D. **(D)** Pictures of two *9x lac* mutants and WT displaying the surface and median views, as well as the overlay of cell wall and fluorescent lignin in the median view. *9x lac, lac1*;*3*;*5*;*7*;*8*;*9*;*12*;*13*;*16*. Lignin was stained with basic fuchsin in **C** and **D** and cell wall was counterstained with calcofluor white M2R in **D**, following a modified version of the Clearsee protocol (41,66). Representative picture at early root development taken at the 15th endodermal cell after onset of endodermal elongation. Scale bars, 20 μm in **B** and 5 μm in **D**. Figure **C** adapted from (40).

### A *per3 9 39 72 64* quintuple mutant is devoid of Casparian strips

Our inability to observe any indication for a role of laccases in CS lignification made it even more pressing to investigate whether peroxidases are indeed of critical importance for CS formation. As mentioned above, strong defects in lignification are observed only upon inhibitor treatments or pharmacological and genetic interference with ROS production, which implicates peroxidase activity only indirectly. Currently, direct genetic manipulation of peroxidases activities only led to partial, quantitative reductions in lignin content (20,36,37). This is almost certainly due to the sizeable number of secreted (type III) peroxidases, with 73 *PER* members in *A. thaliana* (38). Previously, our group had nevertheless attempted to genetically implicate peroxidases in CS lignification in the endodermis, focusing on *PER3* (AT1G05260), *PER9* (AT1G44970), *PER39* (AT4G11290), *PER64* (AT5G42180) and *PER72* (AT5G66390) as those display the highest enrichment of expression in the endodermis (2). As for the laccases in this study, gene expression was confirmed by transcriptional reporters and demonstration that tagged peroxidases are secreted and accumulate at the CS in the endodermis (2) (Fig. S6 and S9A,C). The genetic analysis at the time revealed that a quadruple *T-DNA* insertion mutant of *per3, 9, 39* and *72* (hereafter referred to as *4x per*) did not have any discernable defect in the CS. With no T-DNA insertion lines available for the highly-expressed *PER64* gene, a gene silencing approach was used. By generating an endodermis-specific, inducible artificial micro RNA (amiRNA) against *PER64* in wild-type, a significant delay in endodermal barrier formation could be observed (2). However, it could not be determined whether this partial defect resulted from specific *PER64* silencing, or whether more *PERs* genes were affected by endodermal amiRNA expression. Also, a direct visualization of CS lignification defects was not done at the time.

Using CRISPR-Cas9, we therefore generated both, a single *per64* knock-out mutant in wild-type plants, as well as a quintuple *per3 9 39 72 64* mutant (hereafter designated *5x per*) by transformation into the *4x per* mutant (Fig. S7). We found that the CS of *4x per* plants had no visible lignification defect, appearing identical to wild-type control (Fig. 3). Interestingly, although the *per64* single mutant also displayed a seemingly normal CS, we observed an ectopic deposition of lignin in the edges of the cortex-facing cell wall of the endodermis. Yet, this signal was weak and mostly observed in the surface views (Fig. 3B). More strikingly, we found *5x per* mutant lines to be completely devoid of a CS (Fig. 3B). Interestingly, the endodermis retained its capacity to lignify, as we observed clear ectopic lignin depositions at the corner/edges of the pericycle- and cortex-facing cell walls (Fig. 3A,B), with the intensity of fluorescent lignin at the cortex-facing cell wall being consistently more prominent than the one at the pericycle-facing one. This pattern of ectopic lignification is precisely re-iterating the compensatory lignification observed in many other CS-defective mutants (31,33,39,40). It is known to depend on SCHENGEN (SGN) pathway activation and eventually leads to an alternative, though significantly delayed, formation of an apoplastic diffusion barrier in CS mutants. We therefore assayed diffusion barrier formation in the *5x per* mutants. In wild-type plants, onset of propidium iodide (PI) uptake block occurred at around 14 cells (Fig. 3D) and both single *per64* and the *4x per* mutant showed a similar onset of PI block. By contrast, *5x per* mutant lines displayed a strong delay in apoplastic barrier establishment (Fig. 3D), observed only after about 85 cells in both alleles (Fig. 3D). This delay is much stronger than in other mutants with defective CS and activated SGN pathway, such as *myb36, esb1* or *rbohf* (2,33,39) (Fig. S8). This is explained if the five mutated peroxidases were not only required for CS lignification, but were also partially required for SGN-dependent, compensatory lignification. This would then cause an additional delay in the establishment of the compensatory extracellular barrier through reduced cell corner lignification.

**Figure 3.**
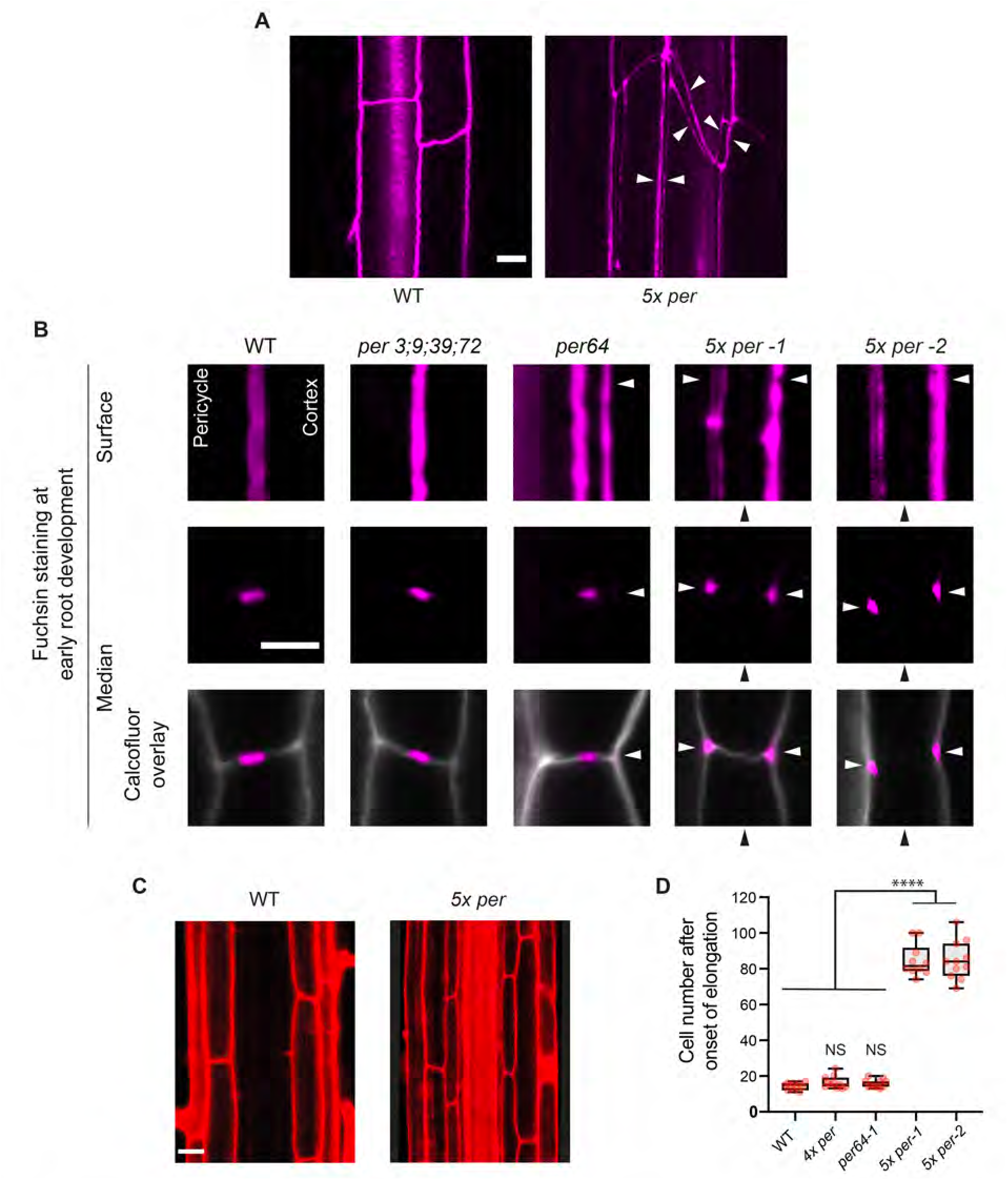
Quintuple peroxidase mutants *per3;9;39;64;72* (*5x per*) are impaired in CS deposition. **(A)** Overview of lignin deposition in the endodermis of plants of wild-type Col-0 (WT) and the *5x per* mutants. White arrows point to ectopic basic fuchsin-positive deposits (lignin-like material). Note that this “double band” appearing in the *5x per* mutant, is cell corner lignification, more easily observed in **B. (B)** Pattern of fluorescent lignin deposition in four peroxidase mutant genotypes compared to WT at early root development (before metaxylem lignification). Pictures displaying the surface and median confocal views, as well as the overlay of cell wall and fluorescent lignin in the median view. *4x per, per3;9;39;72.* Black arrowheads point to the central cell wall position where the CS would normally form. White arrowheads point to ectopic deposits of lignin-like material. Lignin was stained with basic fuchsin and cell wall was counterstained with calcofluor white M2R following a modified version of the Clearsee protocol (41,66). **(C)** The endodermis of *5x per* plants is permeable to the fluorescent apoplastic tracer propidium iodide (red), which enters to the stele whereas in WT it is blocked at the endodermis. Pictures were taken at 20 cells after onset of elongation of endodermal cells. **(D)** *5x per* plants have delayed establishment of apoplastic barrier. Apoplastic barrier function of the endodermis is tested by the propidium iodide diffusion assay. Numbers indicate the endodermal cell in which apoplastic block is established, counting from the onset of endodermis cell elongation (mean, n=10 seedlings per condition; one-way ANOVA, and Tukey’s test as post hoc analysis; ****P<0.0001; NS, not significant). Scale bars, 10 μm in **A;** 5 μm in **B** and 20 μmin **C**.

### A PER64-mCherry fusion construct is able to complement the *5x per* mutant phenotype

We then attempted to complement the phenotype of *5x per* mutants by introducing a C-terminal, fluorescently tagged PER64 variant (*pPER64::PER64-mCherry-4G*) (Fig. S9). For co-visualization of lignin and PER64-mCherry-4G, lignin was stained with auramine-O in this case (41,42), because its green-spectrum emission does not overlap with mCherry. Two independent transformants displayed a CS indistinguishable from wild-type (Fig. S9 and S10), one of which with only some weak, ectopic lignification in some endodermal cells, similar in degree to what is observed in *per64* single mutant (Fig. S10). These slight remaining defects could be explained by either a lower activity of the PER64-mCherry fusion protein or slightly abnormal expression levels of PER64 by the promoter fragment used. Nevertheless, the complementation of the *5x per* mutant by the single PER64-mCherry remains impressive, with one line complementing barrier formation to wild-type levels, while a second line also largely complemented, but with some delay of barrier formation compared to wild-type or the *4x per* mutant (Fig. 3). This also demonstrates that tagging with fluorescent proteins allows PERs involved in lignification to retain their functionality.

### *5x per* plants display strongly increased H_2_O_2_ accumulation in the endodermis

Peroxidases uses H_2_O_2_ as a co-substrate during the oxidation of mono-lignols in the cell wall. We have previously demonstrated that ROS production observed in the endodermal cell wall is fully dependent on a pair of NADPH oxidases, RBOHD and F (2,6). We therefore wondered whether an absence of the peroxidases utilizing ROS for lignin production would impact overall ROS levels. We resorted to the Cerium chloride assay to detect hydrogen peroxide (43). Cerium chloride reacts with H_2_O_2_ to form Cerium perhydroxide, producing electron-dense precipitates, readily detected in transmission electron microscopy (TEM) providing exquisite spatial resolution of ROS detection. This technique has been repeatedly used by our group to describe the pattern of H_2_O_2_ localization in the endodermal cell wall (2,6).

Roots of wild type, *per64* and *4x per* displayed peroxide precipitates that were largely restricted to the outer edge of the CS. Roots of *5x per* lines by contrast, showed very prominent peroxide precipitates along an extended intercellular region between endodermal cells, while no indication of CS formation is observed in this region (Figure 4). This observation perfectly fits the expected consequences of impaired peroxidase activity in endodermal cell walls, which fails to consume the ROS produced by RBOHD and F by polymerizing CS lignin. It corroborates the model postulated for localized lignin deposition in the CS, in which local ROS production by NADPH oxidases is directly used by co-localizing cell wall peroxidases (2). Moreover, it suggests that peroxidase-mediated lignin polymerization represents a powerful ROS-sink *in vivo*.

**Figure 4.**
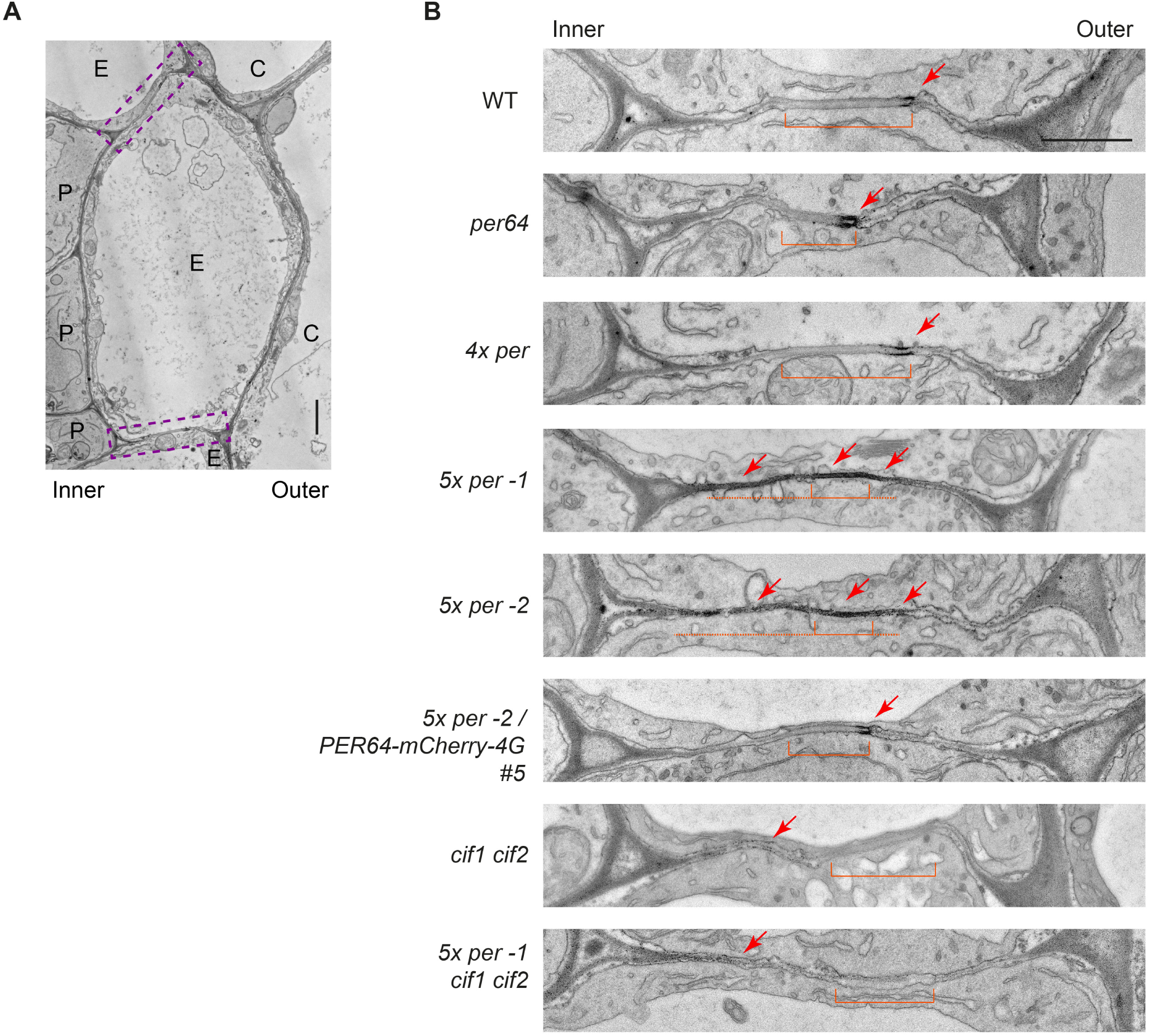
Mutation of *PER 3, 9, 39, 64*, and *72* leads to enhanced and ectopic H_2_O_2_ accumulation in the cell wall around the region where the Casparian strip normally forms. **(A)** Transmission electron micrograph of *A. thaliana* root in cross section. Overview of the endodermal cell wall region displayed in **B**, boxed in purple, dotted line. “Inner” designates the pericycle-facing cell wall, while “Outer” indicates the cortex-facing one. E, endodermis; C, cortex; P, pericycle. WT, wild-type Col-0. **(B)** Micrographs show the median cell wall between endodermal cells. The cells shown correspond to the 1.3 - 1.5 mm region from the root tip of 5-day-old seedlings. Localized H_2_O_2_ production is detected by the cerium chloride (CeCl_3_) assay. H_2_O_2_ reacts with CeCl_3_ to form electron-dense precipitates of cerium perhydroxide. Arrows point to electron-dense precipitates. Brackets, Casparian strip (WT, *per64* and *cif1 cif2*) or the region where the CS would normally localize (5x *per −1, 5x per −2* and *5x per −1 cif1 cif2*). *4x per* carries mutations in *per3, 9, 39* and *72*, and *5x per* carries in addition a mutation in *per64.* Dashed line, extended region of ectopic H_2_O_2_ precipitates. Note absence of cerium perhydroxide precipitates in the median cell wall region in lines carrying the *cif1 cif2* mutations. Scale bars, 1 μm in **A** and **B.**

### Remaining lignification in the endodermis of *5x per* plants is abrogated in SCHENGEN pathway mutants

It was shown that activation of the SGN pathway depends upon the perception of two small signaling peptides, the Casparian strip Integrity Factors 1 and 2 (CIF1 and CIF2) by their receptor, the leucine-rich repeat receptor-like kinase (LRR-RLK) SCHENGEN 3 (SGN3), which in turn activates a receptor-like cytoplasmic kinase, SCHENGEN 1 (SGN1) (4–6). While the CIF peptides are produced in inner cells of the vasculature, SGN1 is localized exclusively to the cortex-facing part of the plasma membrane), outside of the Casparian strip domain (CSD) (6,44). This spatial disposition will lead to continuous activation of the pathway only when CS formation is defective and the diffusion barrier cannot be established. The CIF peptides thus effectively act as diffusion probes, initiating compensatory lignification whenever they succeed in reaching a SGN3-SGN1 signaling complex (6). Recently, it has been demonstrated that SGN3 drives lignification through a branched pathway (6). In one branch, SGN1 directly phosphorylates and activates NADPH oxidases at the plasma membrane, strongly activating ROS production. In the other branch, MAP kinases are activated and an induction of a large set of downstream genes is observed. Two peroxidases, PER15 and 49 are among the most highly upregulated genes upon CIF stimulation ((6) and Fig. S1E). The combination of enhanced ROS production and activation of peroxidase gene expression can directly account for the enhanced and ectopic lignification in cell corners in CS-defective mutants. In order to demonstrate that the ectopic lignification in the *5x per* mutant is also caused by SCHENGEN pathway activation, we generated a *cif1 cif2* double mutant in the *5x per* background, using a specific sgRNA shown to efficiently target both genes (6) (Fig. S11). Plants of *cif1 cif2* have a normally positioned, but discontinuous CS without ectopic lignification (Figure 5). This incomplete CS closure in SGN pathway mutants is thought to reflect a partial requirement for this pathway in undisturbed, wild-type CS formation, possibly to provide a transient boost of CS formation, while the CS is in the process of being built and the diffusion barrier is still permeable. We found *5x per cif1 cif2* plants to be indeed devoid of ectopic lignin in the endodermis in the root zone before metaxylem lignification (Fig. 5). Ectopic lignification could be induced by external application of CIF2 peptide, illustrating that the observed absence of lignification is due to lack of SCHENGEN pathway activation (Figure 5). For reasons we do not understand, a weak, but clearly detectable lignification still seems to form later during root development in the *5x per cif1 cif2* mutant (Fig. S12).

**Figure 5.**
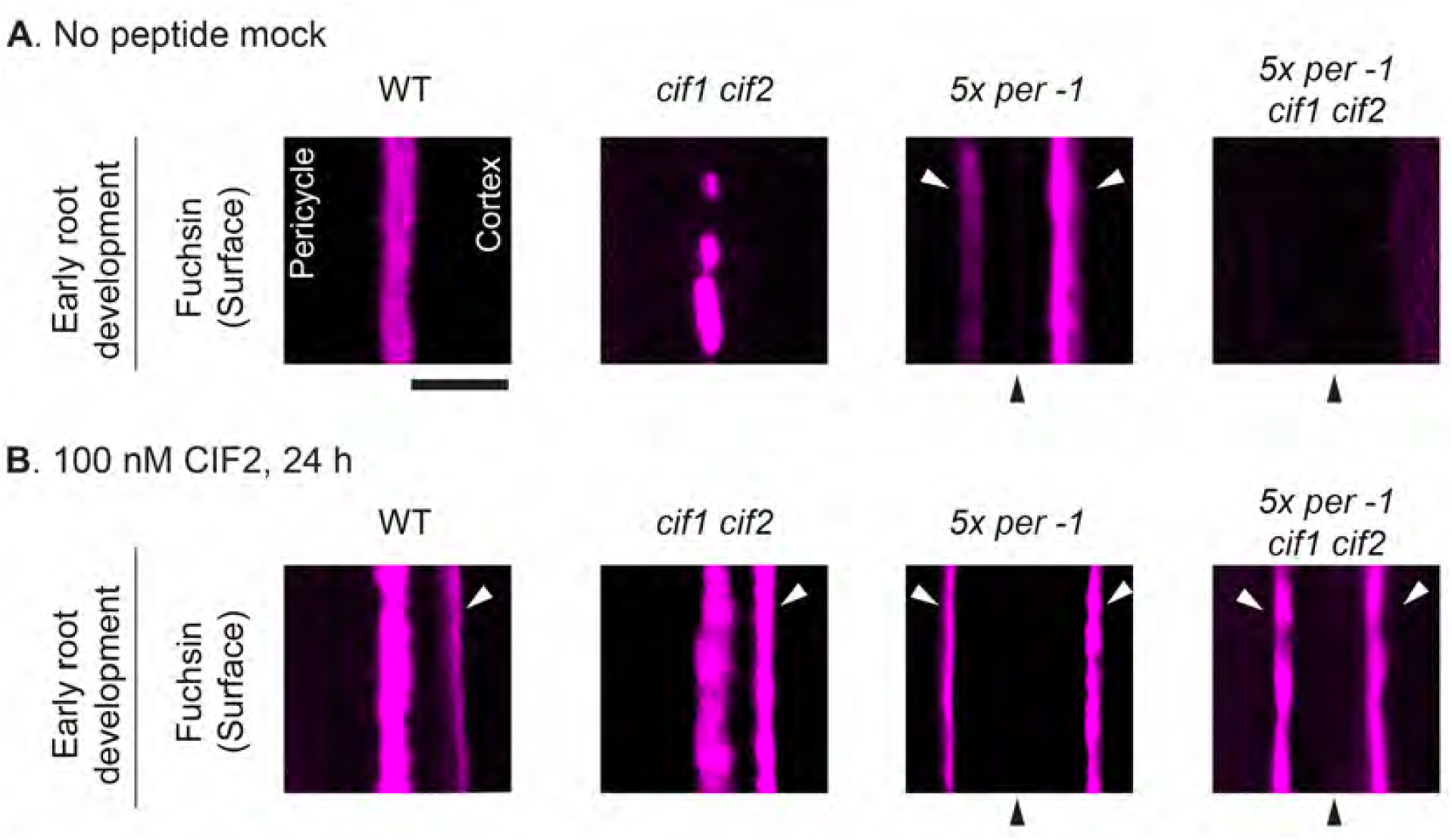
Ectopic deposition of lignin-like material in *5x per* (*3;9;39;64;72*) plants is dependent on the SCHENGEN pathway. **(A)** Surface confocal sections of the Casparian strip (CS) stained with basic fuchsin. Pictures were taken at early root development (before metaxylem lignification). WT, wild-type Col-0. **(B)** Same type of pictures were taken in roots of seedlings treated with 100 nM of SGN3 receptor ligand CIF2 for 24 h. Black arrowheads indicate the wall position where the CS usually forms. White arrows point to ectopic basic fuchsin-positive deposits (lignin-like material). Fluorescent lignin was stained with basic fuchsin and cell wall was counterstained with calcofluor white M2R following a modified version of the Clearsee protocol (41,66). Scale bar, 5 μm.

We also tested whether combining the *cif1 cif2* and *5x per* mutations would further enhance their defects in the establishment of an apoplastic barrier. In the PI diffusion assay *cif1 cif2* and *5x per* mutants have similar delays (barrier closes at 86 ± 9 and 85 ± 9, respectively (Fig. S8B). Nevertheless, *5x per cif1 cif2* plants have a stronger delay than both individual lines, blocking PI after 117 ± 8 cells, essentially encompassing the entire root till the beginning of the hypocotyl (Fig. S8B). This suggests that, while very inefficient, the SCHENGEN signaling pathway still contributes to the very late sealing of the apoplast at the endodermis in *5x per* plants. Interestingly, loss-of-function mutants plants of MYB36, a master regulator of a gene set essential for CS formation, and EXO70A1, a subunit of the exocyst complex required for protein positioning at the CS domain, despite not making any CS, seal the apoplast much earlier than *5x per* plants (39,40) (comparative data shown for *myb36-2* in Fig. S8A). Thus, it seems that mutation of *per3, 9, 39, 64* and *72* not only affects the formation of CS, but also lowers the plant’s capacity to seal the apoplast in the absence of CS by compensatory lignification. This idea is supported by the fact that both, PER9 and 72 (knocked-out in the *5x per* mutant) are significantly induced upon SCHENGEN pathway stimulation ((6) and Fig. S1E).

### The Casparian strip membrane domain (CSD) forms in the absence of lignin and Casparian strip peroxidases

Finally, we checked whether inability to form CS in *5x per* plants was due to a disrupted CSD. We used CASP1-GFP as a marker since this transmembrane protein accumulates very strong and stably at the site of CS formation and is important for its formation. With the exception of *rbohF*, all mutants with a defective CS show a lack or mislocalization of CASP1-GFP (2,40,44–46). We found that CASP1-GFP localizes to the median region of the endodermal plasma membrane, as in wild-type (Figure 6). In addition, ESB1, a secreted protein that strictly co-localizes with CASP1-GFP and is necessary for its correct accumulation and localization, also localizes normally in the cell wall in *5x per* plants (Figure 6). LAC3 also continued to be localized to the CSD (Figure 6). Thus, peroxidases require the CSD for their localization, but the CSD can form independently of their presence and is also unaffected by the absence of lignin.

**Figure 6.**
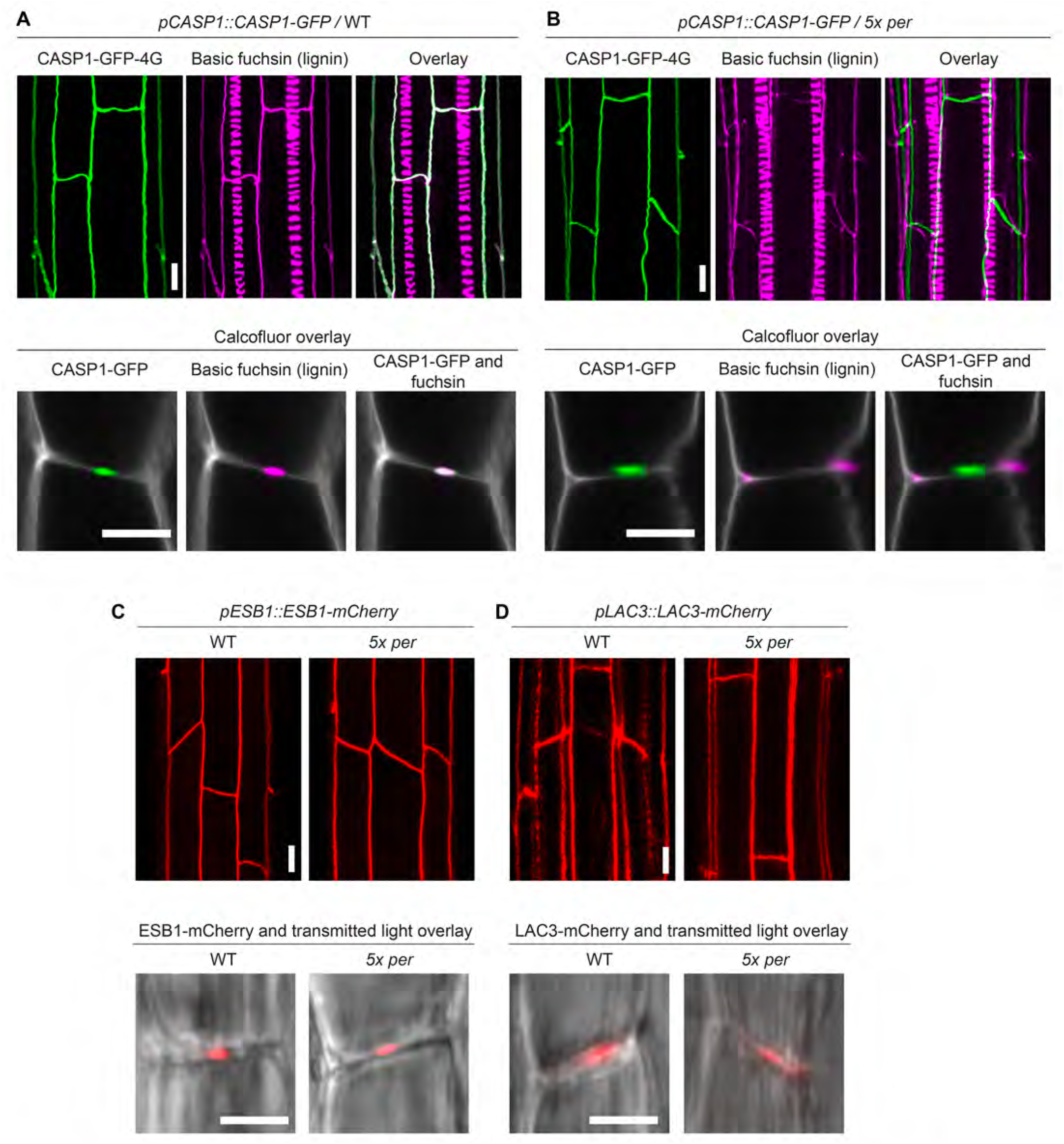
Organization of proteins at the Casparian strip (CS) and CS-membrane domain is unaltered in *5x per* plants. **(A** and **B)** CASP1-GFP-4G (green) labels the Casparian strip membrane domain in wild-type Col-0 (WT) plants (line #3-3-1) and co-localizes perfectly with CS-lignin (magenta, co-localization: white). CASP1-GFP is targeted to the same domain in *5x per* plants (line #4-2-2) and does not co-localize with the cell corner lignin of the mutants. The upper panel shows 3D reconstructions (z-axis maximum projections) of CASP1-GFP-4G and fuchsin. The lower panel shows pictures of median confocal views of the endodermis, displaying the overlay of CASP1-GFP-4G and fuchsin with calcofluor white M2R (gray). **(C)** ESB 1-mCherry is targeted exclusively to the CS in WT, which remains unaffected in *5x* per(line #6-1-1). The upper panel shows 30 reconstructions (z-axis maximum projection) of ESB1-mCherry. The lower panel shows median confocal views of the endodermis with ESB1-mCherry in overlay with transmitted light in gray. **(D)** LAC3-mCherry-4G localizes to the CS and the cortex-facing edges of the endodermal cell wall in WT and is not altered in *5x per* (line #4-6). The upper panel shows 30 reconstructions (z-axis maximum projection) of LAC3-mCherry-4G. The lower panel shows median confocal sections of the endodermis with LAC3-mCherry-4G in overlay with transmitted light in gray. Scale bars in upper panels, 10 µm and in lower panels, 5 μm.

## DISCUSSION

As discussed in the introduction, we hold that the endodermis is an important model for studying lignification, complementing in interesting ways the study of xylem vessels and fibers that represent the vast majority of lignin produced in a mature plant. Yet, in addition to this, lignification in the endodermis might also be more representative of a type of lignification that is widespread, yet constitutes only a minor fraction of total lignin, occurring in cells with no, or little, secondary cell wall and in which lignin is deposited in a restricted fashion as part of a developmental program of differentiation, i.e. during peridermal differentiation or abscission zone formation (17,47), or as part of an induced stress response, i.e. during cell wall damage or bacterial invasion (48,49). Indeed, endodermal lignification occurs in the cell walls between two endodermal cells that amount to a total thickness of barely 200 nm (3,31), very different in dimension from lignified secondary walls, which in the case of interfascicular fibers can be tenfold thicker (ca. 2 µm) (50,51). Beyond this, primary and secondary walls also differ in composition and abundance of their cellulose, hemi-cellulose and pectin components (52,53). This raises the question whether both types of cell walls require different ways of lignification, possibly mediated by different sets of enzymes. Indeed, it was recently reported that xylary fibers localize PER64 in a pattern resembling primary cell walls, while a laccase (LAC4) is localized exclusively to secondary cell wall domains (15).

Previous studies using organ- and plant-wide analysis of *per* mutants only showed quantitative reduction in lignin content (19,20,54–56). Nevertheless, whether *PERs* are indeed essential for lignification has remained unproven. With our focused approach on the CS we now demonstrate that in *5x per* mutants, lignin polymerization in this specific structure is abolished, with a concomitant, local increase in ROS accumulation, illustrating a strong and direct requirement of PERs.

The fact that the *5x per* plants are specifically impaired in CS formation, leaving other lignin structures unaffected, further supports the idea that subsets of *PERs* play very specific roles in various cellular contexts. *PER9* and *PER40*, for example, are required for anther and pollen development likely by enabling polymerization of extensin proteins in the cell wall, while PER36 is required for seed coat mucilage extrusion and is proposed to regulate degradation of the outer cell wall (57,58).

One speculation to account for the strong requirement of peroxidases, yet the lack of any phenotype in a *9x lac* mutant, would therefore be to postulate that laccases are expendable for lignification of primary cell walls, but are required for lignification of secondary cell walls. While this is certainly consistent with and supported by the currently available data, it raises the question why a considerable number of non-required laccases would continue to be expressed in the endodermis and, moreover, localize to CS. Monocotyledonous, more rarely dicotyledonous plants, form very thick secondary cell walls in the endodermis after suberization, that can become lignified, but those occur very late in development (59). We are not aware that the much more short-lived endodermis of Arabidopsis would form any significant amounts of secondary walls that could become lignified. In the future, functional characterization of LACs may help us to better understand their role in lignification.

There are several, not mutually exclusive, ways to explain the presence of laccases in the endodermis: (a) Laccases might be required for CS formation under specific conditions that are not encountered in our laboratory environment, but sufficiently occurrent in nature to maintain their presence in CS. (b) A common transcription factor regulatory network might govern expression of lignification genes for many different cell types and condition. Such a transcriptional “lignification module” might be activated both in cells with secondary walls (requiring laccases) or in those with only primary wall (not requiring them). Supporting such an idea, a MYB transcription factor driving lignification was shown to be upregulated in diverse cell types and conditions, such as pathogen attack, cell wall stress or SGN pathway activation (5,60,61). In such a situation, the presence of “unnecessary” laccases might simply reflect the absence of a dedicated transcriptional module regulating expression of lignification genes in the endodermis. (c) Lack of obvious CS phenotypes in engineered lac mutants may result from RNA-based compensation mechanisms (62), which might support CS formation by upregulation of homologous LACs. Recent studies showed that nonsense mutations created with CRISPR-Cas9 may lead to activation of homologous genes that compensate for the activity of the mutated gene (62). To exclude this latter possibility, a careful monitoring of the remaining LAC genes for upregulated expression in the endodermis will be necessary.

## ACKNOWLEDGMENTS

We thank the Central Imaging Facility (CIF) and Electron Microscopy facility (EMF) of the University of Lausanne for expert technical support. We thank Edouard Pesquet and Edward E. Farmer for discussions on the project.

## AUTHOR CONTRIBUTIONS

N.G. conceived the project. N.R.M., K.H., Y.L., R.U., S.F. and N.G. designed the experiments. N.R.M., K.H., Y.L., R.U., S.F., A.E. and D.D.B. performed the experimental work. N.R.M., K.H. and N.G. wrote the manuscript. All authors revised the manuscript and were involved in the discussion of the work.

## MATERIAL AND METHODS

### Plant material and growth conditions

For all experiments, *Arabidopsis thaliana* (ecotype Columbia) was used. The following mutants and transgenic lines were used in this study: *per3;9;39;72* (*4x per*) (SALK_140204, GABI_186D03, SAIL_369_F11, and SAIL_891_H09, respectively)(2); *esb1-1* (63); *myb36-2* (*GK_543B11*) (46); *rbohf-3* (SALK_059888) (2); *lac1* (SALK_207114); *lac5* (SALK_127394); *lac13* (SALK_023935); *lac16* (SALK_064093); *lac1;5;13;16* (*4x lac*) mutant was created by standard crossing procedures. For a list of primers used for mutant genotyping, see Table S1. The following mutants were generated by CRISPR-Cas9 method (34,35). For *per64*, a protospacer sequence compatible with the SpCas9 and StCas9 in (35) were used in independent experiments. The SpCas9-compatible protospacer was deployed in the *4x per* genetic background to create *5x per* mutants. To obtain *lac3* alleles we utilized a single protospacer sequence. LAC3 was mutated in the *4x lac* genetic background to obtain the *5x lac* mutants. In order to generate *9x lac* mutants a multiplex sgRNA cloning strategy was followed to target *LAC7, 8, 9* and *12* in the *5x lac* genetic background. Two sgRNAs were deployed simultaneously per each of these genes, except for *LAC9*, where only one sgRNA was used, which has a common target in *LAC8. cif1 cif2* mutants were generated using a sgRNA that targets both CIF1 and CIF2. A list with the sgRNA sequences is available in Table S2. Protospacer sequences were designed with the CRISPR-P 2.0 design tool (http://crispr.hzau.edu.cn/CRISPR2/) (64) and Benchling (https://benchling.com).Transgenic plants with expression constructs were generated by transformation of *Arabidopsis* plants by the floral dip method (65). Seeds were surface-sterilized by quick rinse in 70% EtOH, followed by 96% EtOH, dried and plated on half-strength MS (Murashige and Skoog medium) containing 0.8% agar plates and vernalized at 4 °C for 2-3 days in the dark. Seedlings were grown vertically at 22 °C, under long-day conditions (18 h, 100 µE). All microscopic analyses were performed on roots of 5-day-old seedlings. For assays with CIF2 peptide, 4-day-old seedlings were transferred to half strength MS plates supplemented with 1µM, 100 nM or water (untreated) for 1 day.

### Plasmid construction

Plasmids constructs were generated by Multisite gateway technology (Thermo Fisher Scientific). Promoter sequences were flanked by KpnI and XmaI sites, cloned into a p4-p1r entry vector, and then recombined by LR reaction to p221 NLS & p2r-p3 GFP-GUS to generate transcriptional reporters. The following promoters were used (bp before ATG): *pLAC1* (1904 bp), *pLAC3* (2111 bp), *pLAC5* (2235 bp), *pLAC7* (2021 bp), *pLAC13* (2060 bp), *pLAC16* (2011).

For translational fusion constructs a destination vector of the type *pDEm34GW,0* was used. The following expression plasmids were generated using the same promoter fragments cited above: *pLAC1::LAC1gDNA-mCherry-4G* (4 glycine extension), *pLAC3::LAC3gDNA-mCherry-4G, pLAC5::LAC5gDNA-mCherry-4G, pLAC13::LAC13gDNA-mCherry-4G, pLAC16::LAC16gDNA-mCherry-4G*. In addition the following translational fusions where created using the *pCASP1* promoter: *pCASP1::LAC1gDNA-mCherry-4G, pCASP1::LAC3gDNA-mCherry-4G, pCASP1::LAC5gDNA-mCherry-4G, pCASP1::LAC13gDNA-mCherry-4G, pCASP1::LAC16gDNA-mCherry-4G.*

Cloning of plasmids used for CRISPR-Cas9 was performed as previously described. Briefly, the spacer to customize sgRNA for SpCas9 or StCas9 was cloned by annealing the oligos, then ligated into *Bbs*I linearized, Gateway-entry plasmid *pEn-Chimera* (34). For multiplex targeting of *LAC7, 8, 9* and *12*, the six sgRNAs utilized were cloned into two vectors, each carrying 3 sgRNAs. *5x lac* plants were co-transformed with both plasmids.

### Histological techniques

The propidium iodide (PI) diffusion assay was performed as previously described. 5-day-old seedlings were incubated in a solution of 15 µM propidium iodide for 10 min in the dark, then incubated in water for 30 seconds, mounted in water using a slide and coverslip and immediately observed in microscope. Block of PI in the endodermis was quantified in terms of the number of endodermal cells after onset of elongation. Onset of elongation was defined as the point where the length of an endodermal cell was more than twice its width. Col-0 was always included as control.

For visualization of Casparian strips, cell wall and fluorescent proteins a modified protocol of ClearSee was used (41,66). The Casparian strip was stained with the lignin dye basic fuchsin (0.2%. Fluka, Analytical, CAS-No: 58969-01-0). Polysaccharide cell wall was stained with 0.1% calcofluor white M2R (Polysciences, CAT#4359).

### Confocal laser scanning microscopy (CLSM) imaging

Confocal laser scanning microscopy experiments were performed on a Zeiss LSM 880 or a Leica SP8X microscope. All combinatorial fluorescence analyses were run as sequential scans. The following excitation and emission settings were used to obtain specific fluorescence signals: propidium iodide, 488 nm/600–620 nm; Basic Fuchsin, 561 nm/570–650 nm; Auramine-O, 488 nm/505-530 nm; EGFP, 488 nm/ 500–550 nm; mCherry, 561 nm/ 600–650 nm; Calcofluor white, 405 nm, 425–475 nm. Confocal images were processed and analyzed using the Fiji package of ImageJ (http://fiji.sc/Fiji).

### Transmission electron microscopy for detection of H_2_O_2_ by the Cerium chloride assay

Visualization of H_2_O_2_ around the Casparian strip was done by cerium chloride method as described in previous reports (2,6,43) with some modifications. The histochemical method is based on the generation of cerium perhydroxides as described and was used for the location of H_2_O_2_ at the Casparian strip. Cerous ions (Ce^3+^) react with H_2_O_2_ forming electron-dense cerium perhydroxide precipitates, which are detected by electron microscopy.

Five-day-old seedlings were incubated in 50 mM MOPS (3-[N-morpholino] propane sulphonic acid) (pH 7.2) containing freshly prepared 10 mM cerium chloride (CeCl_3_) for 30 min. After the incubation with CeCl_3_, seedlings were washed twice in MOPS buffer for 5 min and fixed in glutaraldehyde solution (EMS, Hatfield, PA) 2.5% in 100 mM phosphate buffer (pH 7.4) for 1 hour at room temperature. Then, they were post-fixed in osmium tetroxide 1% (EMS) with 1.5% of potassium ferrocyanide (Sigma, St. Louis, MO) in phosphate buffer for 1 hour at room temperature. Following that, the plants were rinsed twice in distilled water and dehydrated in ethanol solution (Sigma) at gradient concentrations (30%, 40 min; 50%, 40 min; 70%, 40 min; 100%, 1 h, twice). This was followed by infiltration in Spurr resin (EMS) at gradient concentrations (Spurr 33% in ethanol, 4 h; Spurr 66% in ethanol, 4 h; Spurr 100%, 8 hours, twice) and finally polymerized for 48 hours at 60°C in an oven. Ultrathin sections 50 nm thick were cut transversally at 1.3 ± 0.1 mm from the root tip, on a Leica Ultracut (Leica Mikrosysteme GmbH, Vienna, Austria) and picked up on a nickel slot grid 2×1 mm (EMS) coated with a polystyrene film (Sigma). Micrographs were taken with a transmission electron microscope FEI CM100 (FEI, Eindhoven, The Netherlands) at an acceleration voltage of 80kV and 11000 magnifications (pixel size of 1.851nm, panoramic of 17 × 17 pictures), exposure time of 800ms, with a TVIPS TemCamF416 digital camera (TVIPS GmbH, Gauting, Germany) using the software EM-MENU 4.0 (TVIPS GmbH, Gauting, Germany). All the pictures were taken using the same beam intensity, and panoramic aligned with the software IMOD (67).

---

**Supplementary figure 1.**
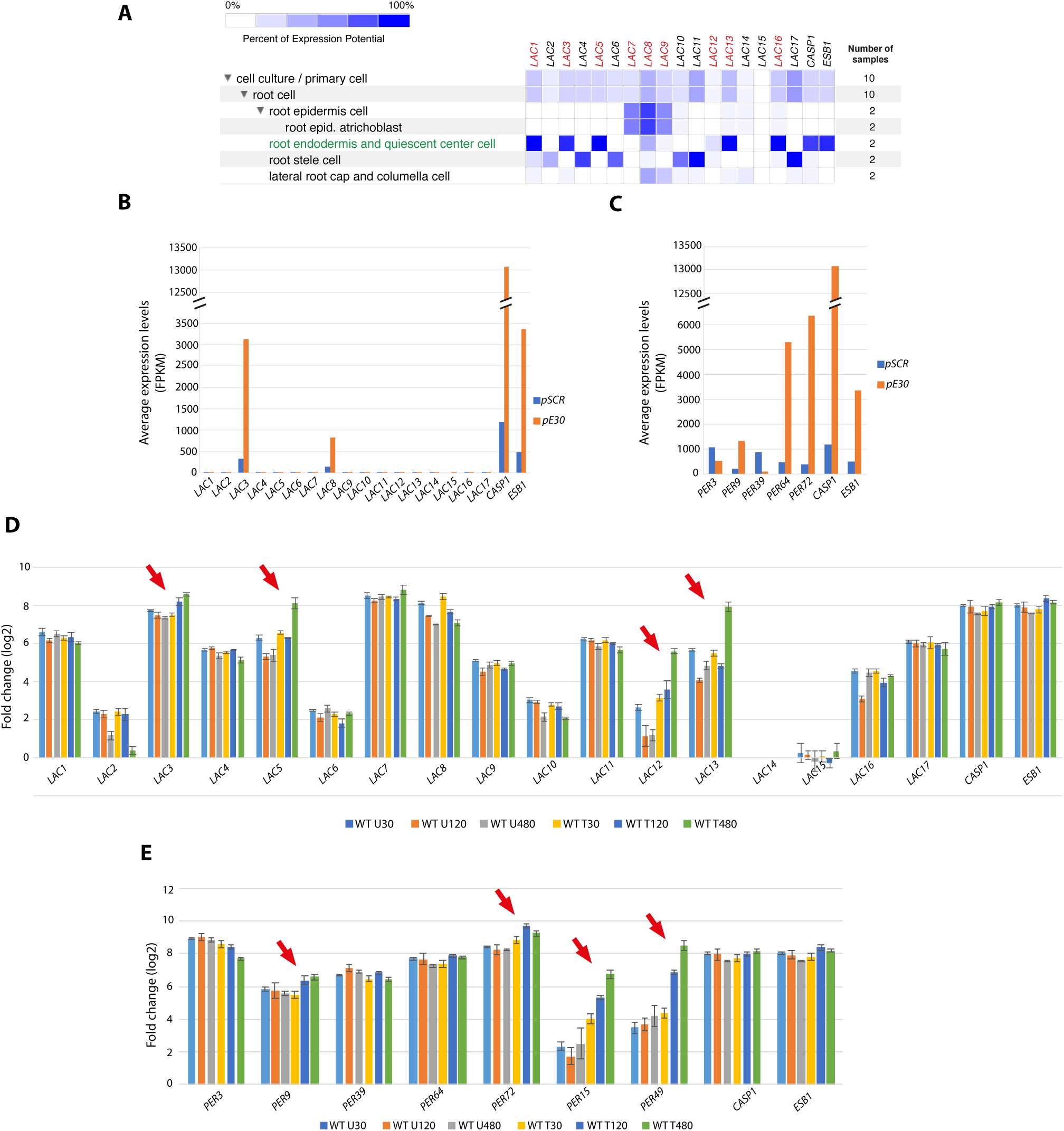
Database expression analysis of *LAC* and *PER* genes in root tissues of *A. thaliana* seedlings. **(A)** mRNA expression pattern of *LACs* in Arabidopsis screened by Genevestigator (1). Dataset of tissue-specific translatome/RNA sequencing (2). Heat maps represent absolute expression values as shading intensity computed in Genevestigator and plotted in a linear scale. Percent of expression potential: normalized average relative to the top 1 percent values of corresponding genes across all samples included in Genevestigator. Only wild-type genotypes and mock, non-treated samples were considered. *LAC* genes mutated in *5x lac* and *9x lac* lines in this study are labeled in red. Datasets for endo-dermis-specific expression are labeled in green. Data for the endodermis-specific genes *CASP1* (At2g36100) and *ESB1* (At2g28670) are also shown. **(B-C)** Dataset of endodermis-specific transcriptome/RNA sequencing (3). **(B)** Data for the *LAC* family members. **(C)** Data for the *PERs* whose products are known to be targeted to the Casparian strip (2). *pSCR*, early-differentiating endodermis gene promoter. *pE30*, mature endodermis gene promoter. GFP markers driven by those promoters were used to prepare endodermis protoplasts for mRNA-profiling. FPKM, fragments per kilobase pairs per million reads. **(D-E)** Root mRNA profiling after external treatment with the CIF2 small peptide ligand (4). CIF2 activates the SGN pathway leading to enhanced endodermal lignification. mRNA profiling was performed in a time course including 30 min, 120 min and 480 min after CIF2 (WT T) or mock (WT U) treatment. **(D)** Data for the *LAC* family members. **(E)** Data for the *PER* members analyzed in this study as well for *PERs* previously shown to be induced by CIF2 treatment (4). Data for endodermis specific genes *CASP1* and *ESB1* are presented for comparison. Red arrows point genes with induced expression upon CIF2 treatment.

**Supplementary figure 2.**
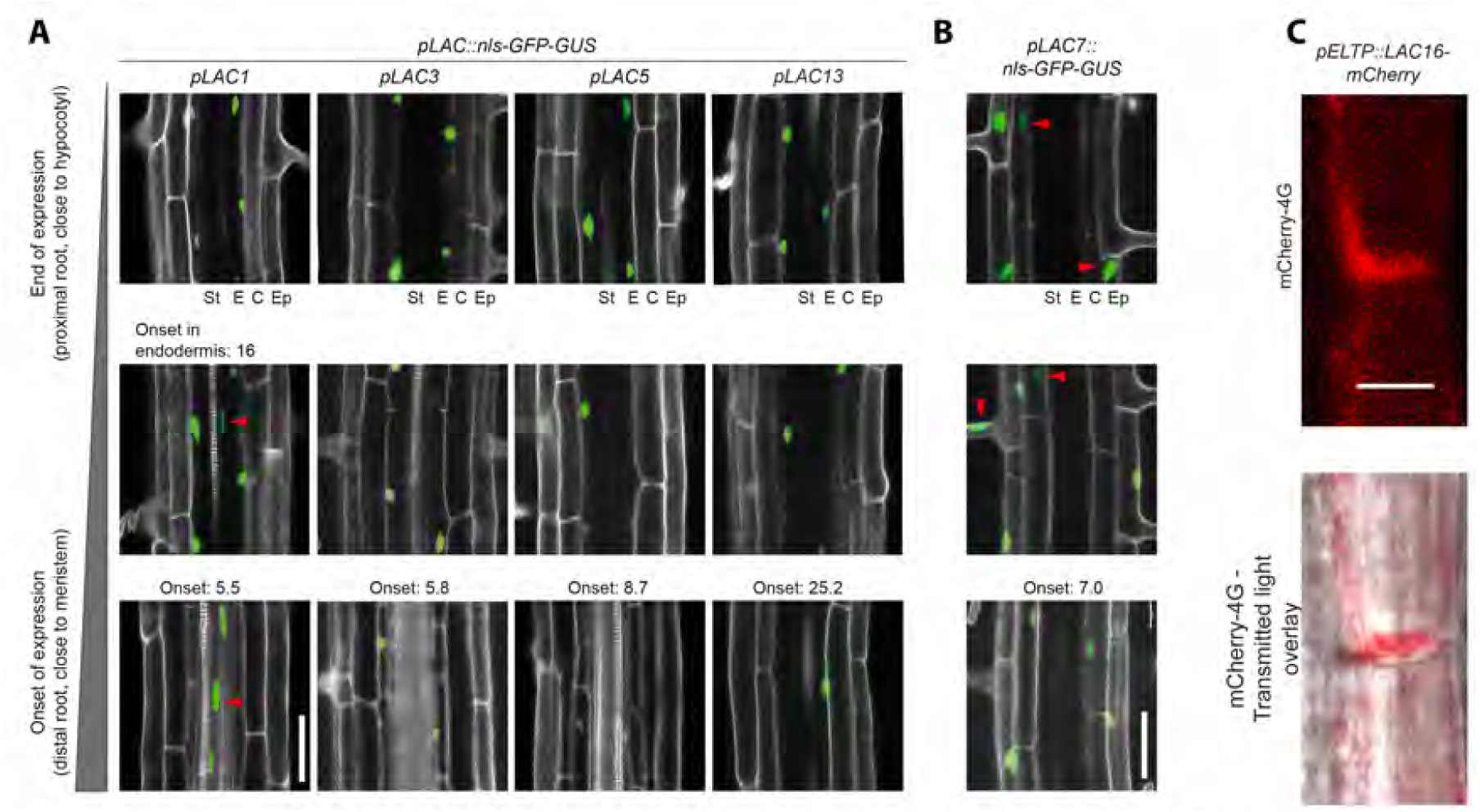
Validation of *LAC* gene expression in the endodermis of *A. thaliana.* **(A)** Tissue specific expression of *pLAC::nls-GFP-GUS* transcriptional reporters along the root of 5-day old seedlings. Pictures show the overlay of GFP (green) and propidium iodide (gray) signals acquired with confocal laser scanning microscopy. Onset, first cell where GFP expression is observed, using as reference the onset of endodermis cell elongation. This reference is used irrespective of the tissue where GFP is expressed. St, stele. E, endodermis. C, cortex. Ep, epidermis. Red arrow points to expression of nls-GFP marker in tissues other than the endodermis. The four pictures in the central panel of this figure are identical to Fig.1A. **(B)** *pLAC7::nls-GFP-GUS* transcriptional reporter was not active in the endodermis. Legends as in **A. (C)** Localization of LAC16-mCherry-4G expressed by the endodermis-specific *ELTPIEDA4* promoter *(pELTP).* The signal pattern of the fusion protein suggests cytoplasmic localization. Living 5-day-old seedlings were imaged by Confocal laser scanning microscopy. Scales bar are 50 µm in **A** and **B**, and 10 μm in **C.**

**Supplementary figure 3.**
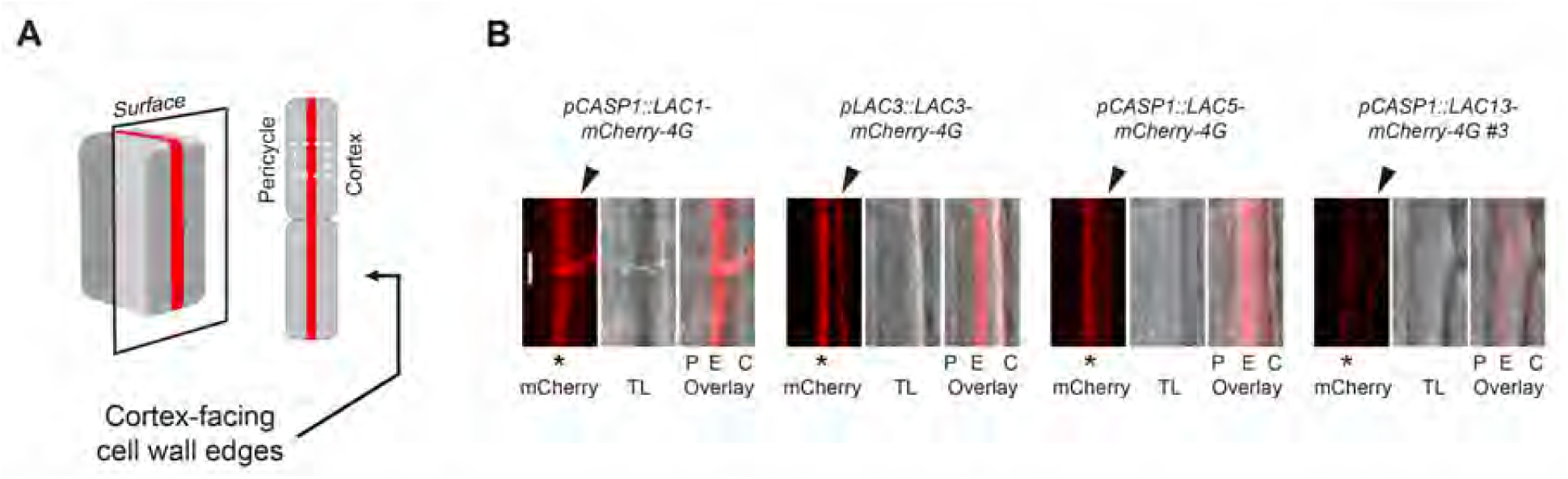
Subcellular localization of LAC1-, LAC3-, LAC5- and LAC13- mCherry-4G to the median position of the endodermis cell wall. **(A)** Schematic of the region of endodermal cells imaged and shown in the pictures in **B.** Schematic adapted from (5). **(B)** C-terminal, fluorescently tagged proteins of the 4 LACs localize to the median position of the endodermal cell wall, where the CS is located, as well as to the outer edges/corners of the endodermal cell wall. Pictures are from surface confocal views, using 5-day-old living seedlings. The gray channel corresponds to transmitted light (TL). P, pericycle. E, endodermis C, cortex. Asterisks indicate median cell wall position, where the CS forms. Black arrowheads point to the cortex-facing edges of the cell wall in the endodermis. Scale bar in **B**, 5 μm.

**Supplementary figure 4.**
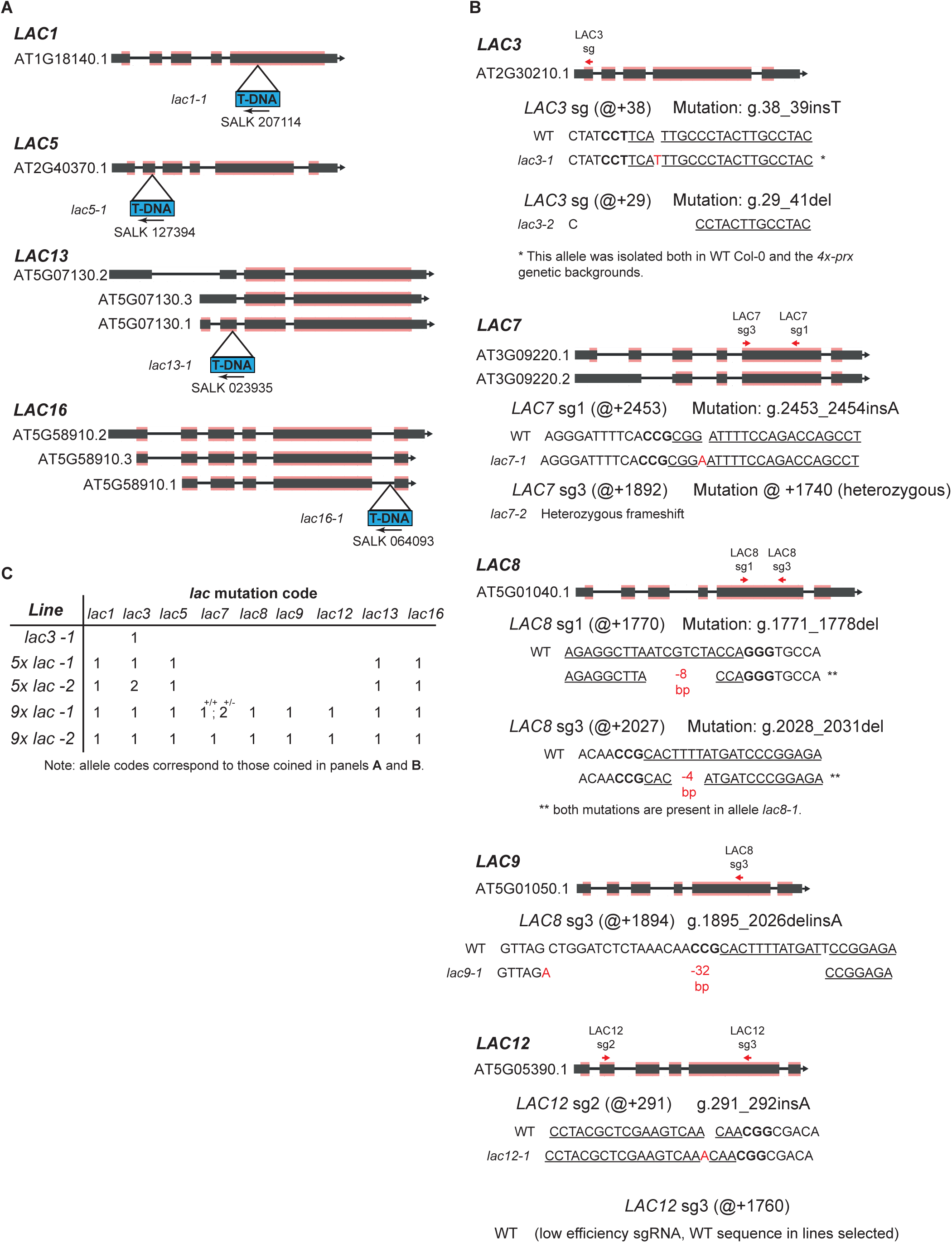
Mutant alleles of *LAC1, 3, 5, 7, 8, 9, 12, 13* and *16* in the respective nonuple mutant. **(A)** Transfer DNA (T-DNA) insertion alleles of *LAC1, 5, 13* and *16*. Dark grey rectangle, transcribed region; dark-grey rectangle with pink layers, coding DNA sequences; lines, introns. Triangles, T-DNA insertion site. Arrows indicate the directionality of the T-DNA left border. Gene AT codes are provided for each gene. **(B)** CRISPR-Cas9 induced mutant alleles of *LAC3, 7, 8, 9* and *12.* Schematic of *LAC* genes and CRISPR spacers utilized. Gene editing was deployed in the quadruple T-DNA *lac1;5;13;16* (*4x lac*) background. Red arrows, localization of target loci. Two red arrows in a gene indicate that two sgRNAs were used simultaneously for mutagenesis. @+ symbol on WT sequence, position of mutation counting from the start codon. Underlined sequence, CRISPR spacer. Bold text, PAM sequence. Red characters in allele sequences indicate gene mutations. Other symbols same as in **A**. Nomenclatural description of mutations based on nucleotide numbering are provided as previously suggested (6). For example, g.38_39insT, indicates a T-insertion (ins) between nucleotide 38 and 39 counting from the gene (g) start codon. In g.29_41del, the mutation is a deletion (del) of nucleotides 29 to 41, counting from the gene start codon. **(C)** Overview of mutant *lac* alleles present in the lines used in this study. Numbers indicate the allele code for every *lac* mutation. +/+, homozygous mutation; +/-, heterozygous mutation.

**Supplementary figure 5.**
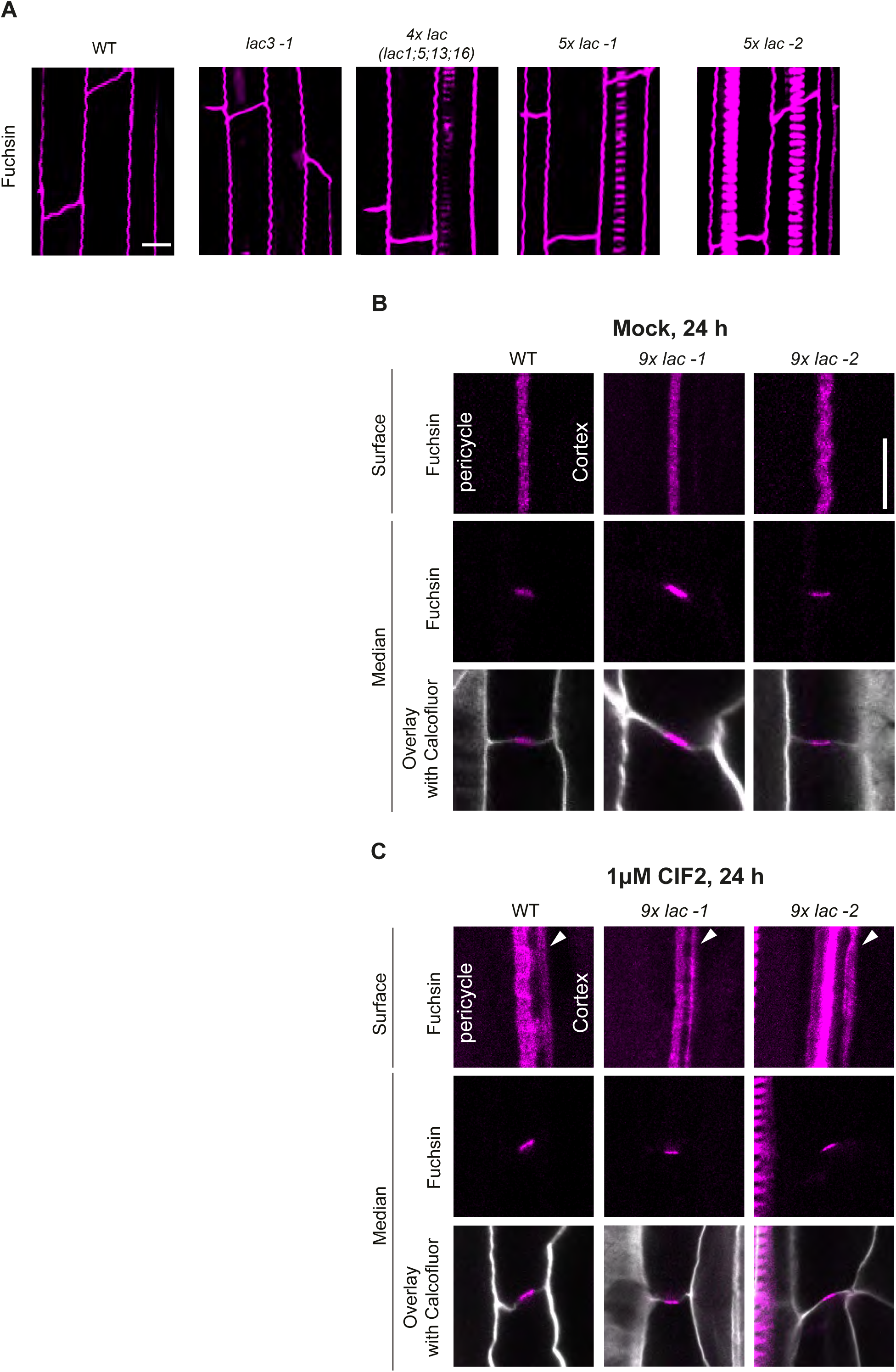
A quintuple and nonuple mutant in *lac1, 3, 5, 7, 8, 9, 12, 13* and *16* have no discernible defect in endodermis lignification. **(A)** *5x lac* mutants (*lac1*;*3*;*5*;*13*;*16*) and *lac3-1* produces continuous CS undistinguishable to the one in the wild-type Col-0 (WT). 3D projection of root samples stained with basic fuchsin. **(B - C)** In *9x lac* (*lac1, 3, 5, 7, 8, 9, 12, 13* and *16*) mutants the endodermis is sensitive to CIF2-induced deposition of lignin-like material. **(B)** mock and **(C)** seedlings treated with 1 μM CIF2 small peptide (dissolved in water) during 24 h. Surface and median confocal views of the CS. White arrows point to ectopic, basic fuchsin-positive deposits (lignin-like material). In **A, B** and **C** fluorescent lignin was stained with basic fuchsin and cell wall was counterstained with calcofluor white M2R following a modified version of the Clearsee protocol (7,8). Scale bar, 5 µm.

**Supplementary figure 6.**
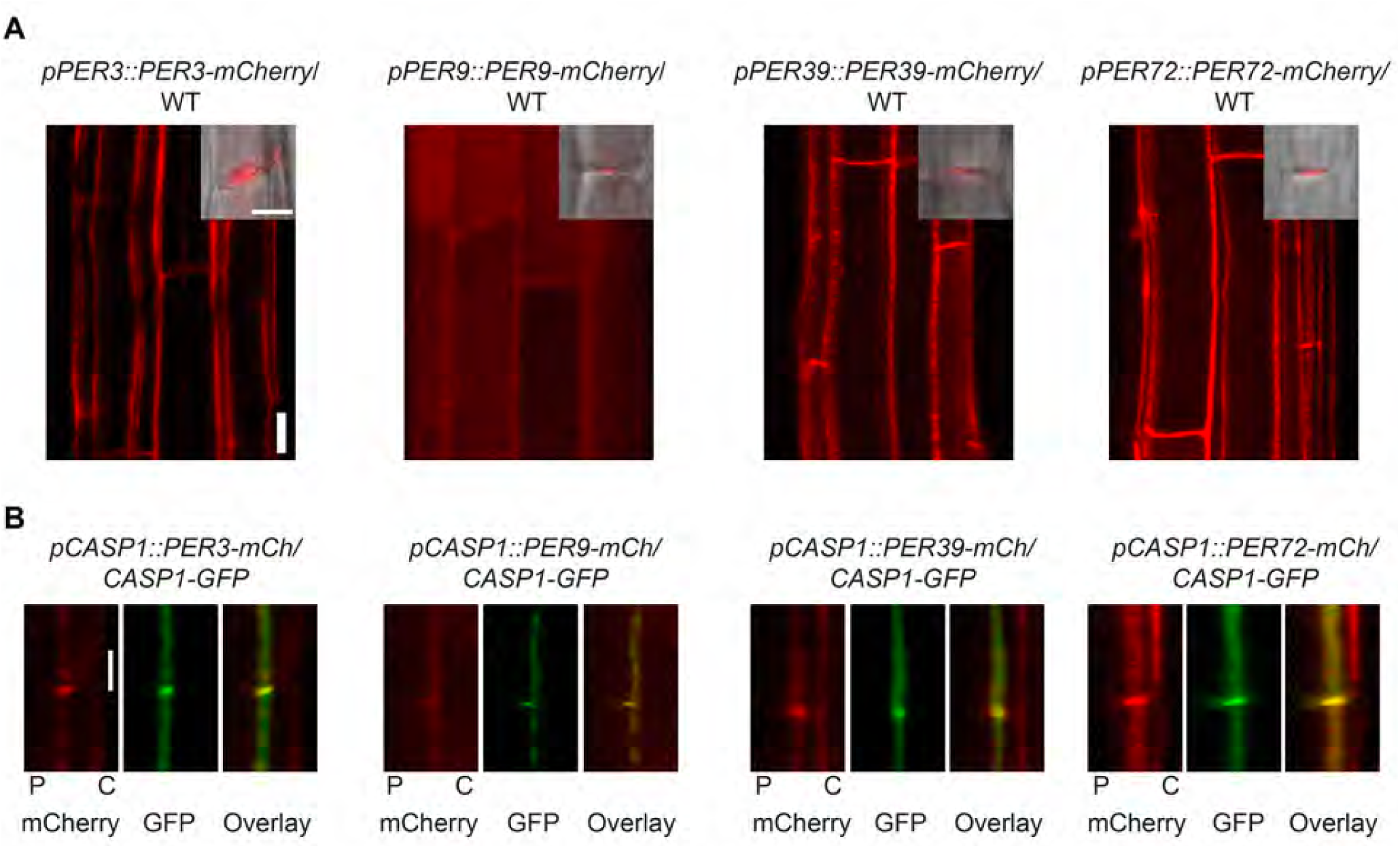
Subcellular localization of PER3-, PER9-, PER39- and PER72- mCherry-4G to the Casparian strip (CS). **(A)** C-terminal, fluorescently tagged proteins of the 4 PERs localize at the CS. In addition, PER 3, 39 and 72 are also observed at the outer edges/corners of the endodermal cell wall. Inset of the median confocal section of the localization of the 4 PERs to the median position of the endodermal cell wall. The gray channel corresponds to transmitted light. WT, wild-type Col-0. **(B)** The 4 PERs co-localize with the Casparian strip-membrane-domain protein CASP1-GFP. Recruitment of the 4 PERs to the CS is first observed in the root zone of CASP1-GFP-expression onset. Localization of PER3, PER39 and PER72 at the CS remains along the root, whereas PER9 is only transiently recruited to the CS in the first 1-2 endodermal cells after onset of CASP1-GFP localization. Median and surface sections of CS are as explained in Figures 1C and S3A. Lines imaged for this figure were originally reported in (9). Localization of PER64 is presented in Figure 4. Scale bars in are 10 μm and 5 μm in the main and inset pictures in **A**, respectively; and 5 μmin **B.**

**Supplementary figure 7.**
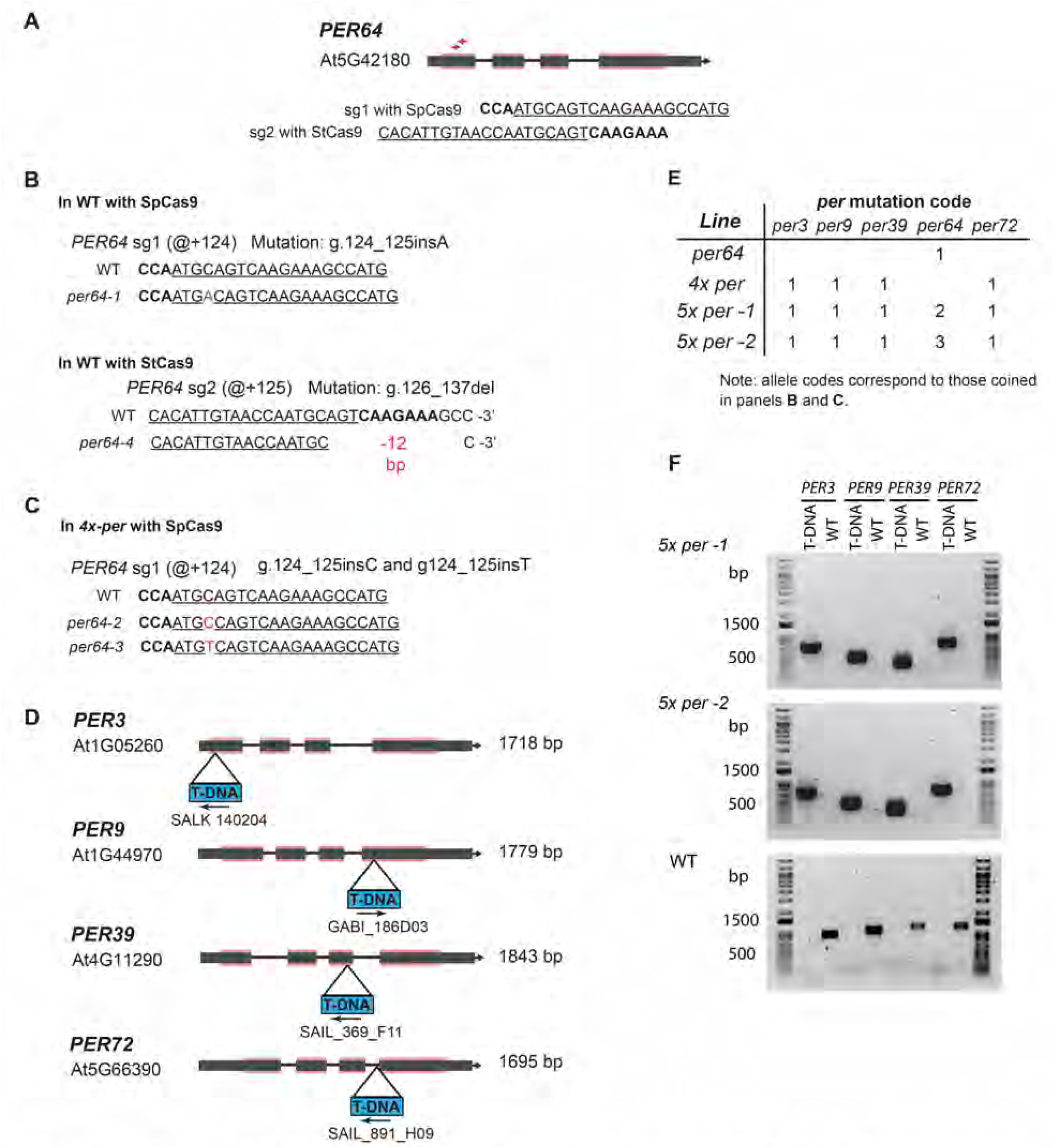
Generation of *per64* single and quintuple *per3; 9; 39; 64; 72* (*5x per*) mutants. **(A)** Schematic of the *PER64* gene and CRISPR spacers (spacers) used in this study. Dark grey rectangle, transcribed region; dark-grey rectangle with pink layers, coding DNA sequences; lines, introns. Two sgRNAs were used separately, one compatible with the *Streptococcus thermophilus* Cas9 nuclease (StCas9) and another with the S. *pyogenes* Cas9 nuclease (SpCas9). These spacers have overlapping targeting regions at the first exon of *PER64.* Red arrows, localization of target loci. Underlined sequence, CRISPR spacer region. Bold text, PAM sequence. **(B), (C)** Mutant alleles of *per64* obtained in wild-type Col-0 (WT) using sgRNAs for SpCas9 and StCas9, and in the quadruple T-DNA mutant *per3;9;39;72* (*4 x per*) genetic backgrounds using the SpCas9 sgRNA. @+ symbol on WT sequence, position of mutation counting from the start codon. Red characters in allele sequences indicate gene mutations. Description of mutation following the nomenclature in (6). **(D)** Transfer DNA (T-DNA) insertions in *PER3, 9, 39* and *72* present in the *4x per* line (9) used in this study. Triangles, T-DNA insertion site. Arrows indicate the directionality of the T-DNA left border. Other symbols same as in **(A).** (**E**) Overview of mutant *per* alleles present in the lines used in this study. Numbers indicate the allele code for every *per* mutation. (**F**) Genotyping by PCR-agarose gel electrophoresis of T-DNA insertions in the two *5x per* lines used in this study.T-DNA, PCR amplicon spanning the left border of T-DNA insertions. WT, PCR amplicon of the T-DNA insertion flanking regions to test for absence of insertion.

**Supplementary figure 8.**
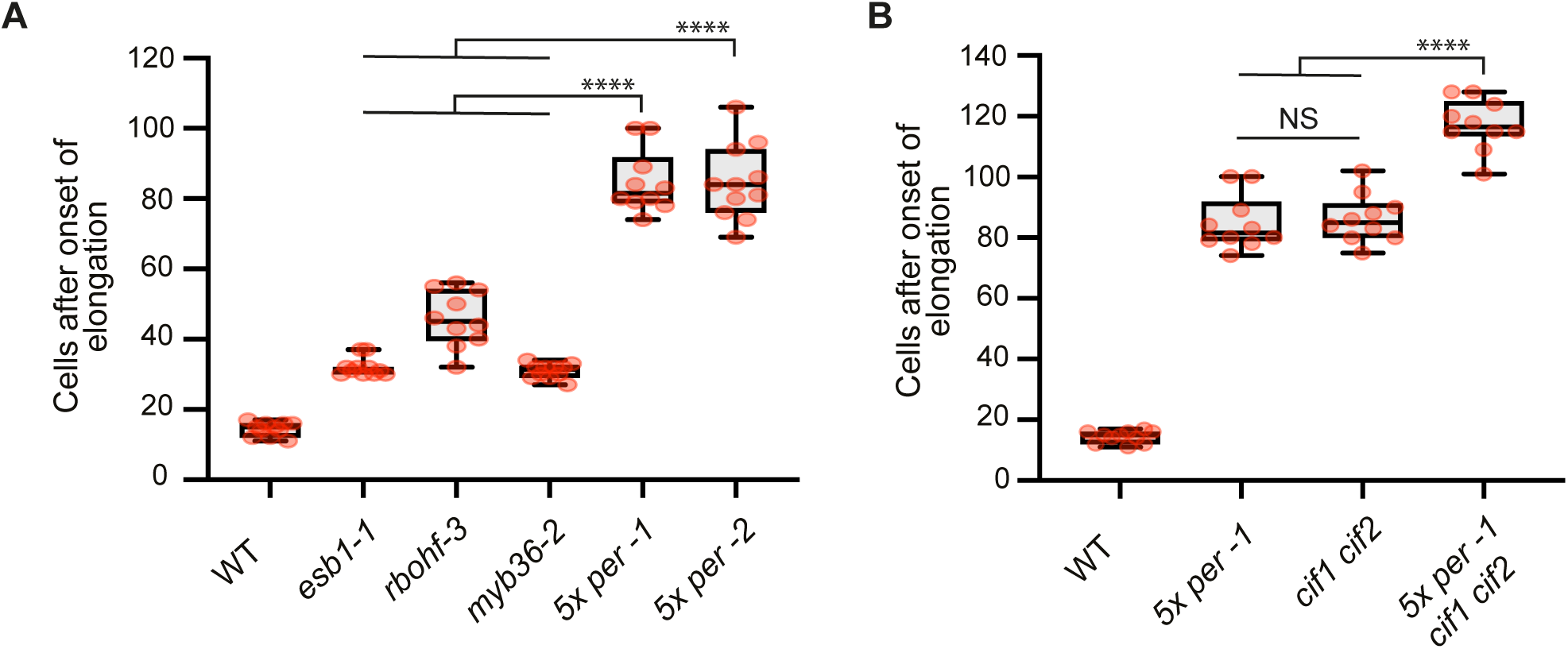
Comparison of delay in apoplastic block establishment between distinct Casparian strip (CS) mutants by PI diffusion assay. **(A)** Comparison of *5x-per* against mutants of *RBOHF* and *ESB1*, genes involved in CS lignification, and *MYB36*, a master transcription factor for CS formation. WT, wild-type Col-0. **(B)** Comparison of *5x-per* against the mutant in *CIF1* and *CIF2*, genes encoding small peptide ligands, which activate the formation of compensatory lignin-like material in the endodermis. Apoplastic barrier function of the endodermis is tested by the propidium iodide diffusion assay in **A** and **B**. Numbers indicate the endodermal cell in which apoplastic block is established, counting from the onset of endodermis cell elongation (mean, n=10 seedlings per condition; one-way ANOVA, and Tukey’s test as post hoc analysis; ****P<0.0001).

**Supplementary figure 9.**
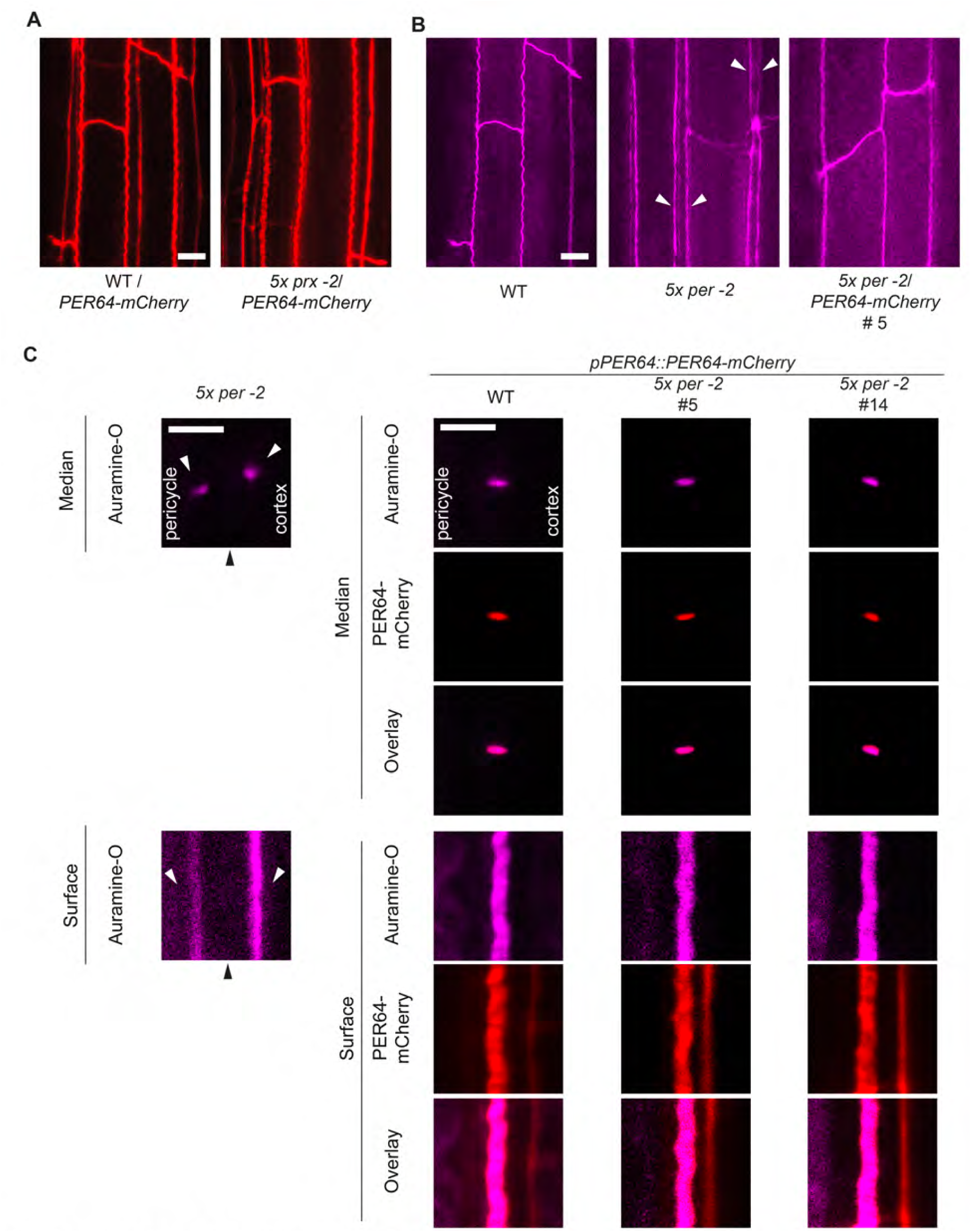
Complementation of Casparian strip absence in *5x per* with a copy of *PER64* (*pPER64::PER64-mCherry-4G*). **(A)** Localization pattern of PER64-mCherry-4G in wild-type Col-0 (WT) and *5x per* complemented line. **(B)** Overview of lignin deposition in the endodermis of plants of Col-0, *5x per* −2 and the mutant complementation line *5x per* −2 *I PER64-mCherry-4G* #5. White arrows point to ectopic auramine-O - positive deposits (lignin-like material). **(C)** Pattern of fluorescent lignin deposition in plants of WT compared to *5x per* −2 and two independent mutant complementation lines, *5x per −2/PER64-mCherry-4G* #5 and #14. Pictures display the surface and median confocal views of fluorescent lignin and PER64-mCherry-4G. Black arrowheads point to the central cell wall position where normally the CS forms. White arrowheads point to ectopic deposits of lignin-like material. Lignin is stained with auramine-O following a modifying version of the Clearsee protocol (7,8). Scale bars, 10 µmin **A, B;** and 5 μmin **C.**

**Supplementary figure 10.**
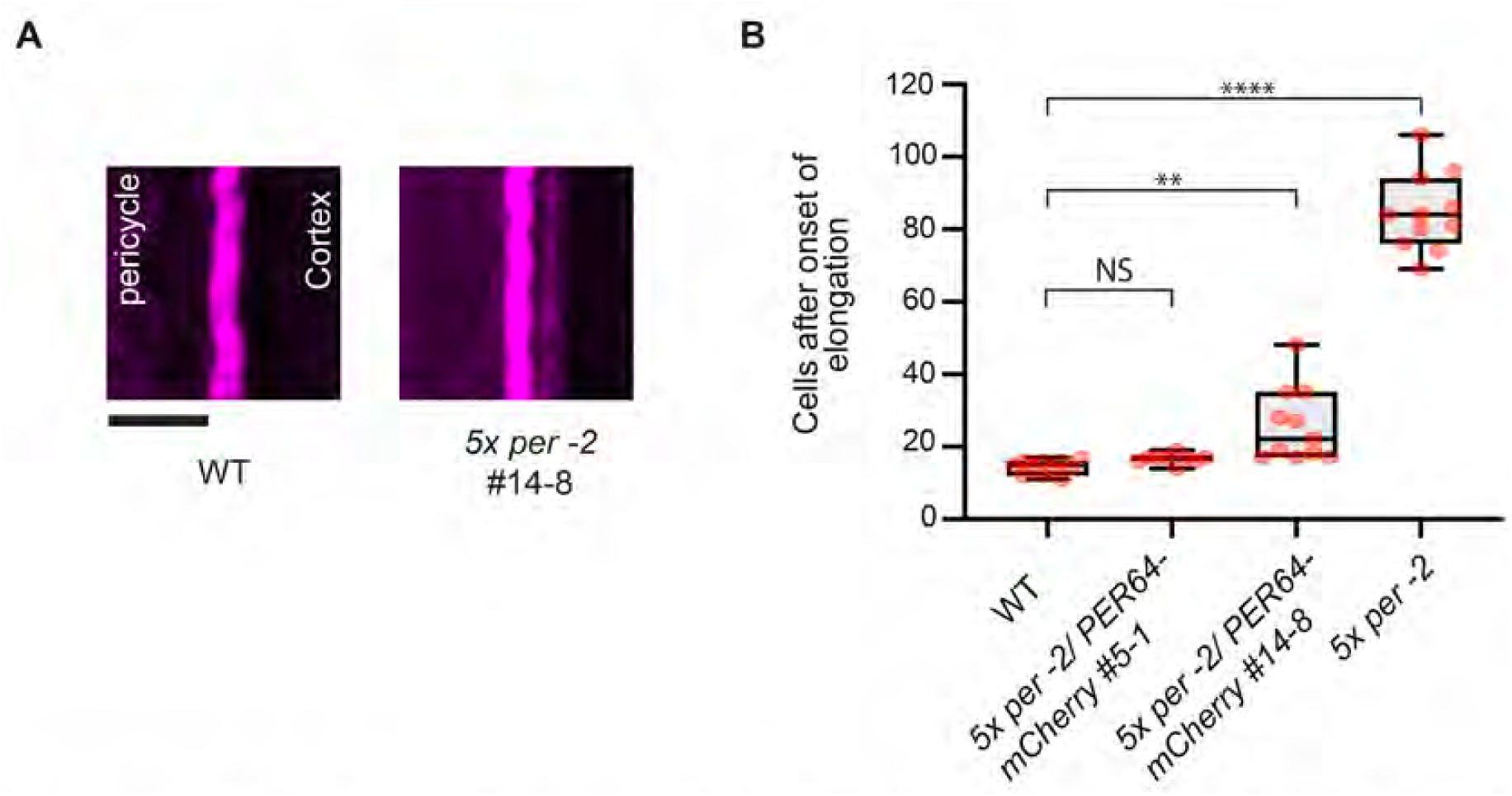
Complementation of the Casparian strip (CS) mutant phenotype in *5x per* plants. **(A)** Some endodermal cells in the mutant complementing line #14-8 have a CS and faint, ectopic auramine-O- positive deposits (lignin-like material) at the cortex-facing cell wall. Pictures display surface confocal views. Lignin is stained with auramine-O following a modified version of the Clearsee protocol (7,8). Additional pictures of line 14-8 are shown in Fig. 4C. Scale bar, 5 μm. WT, wild-type Col-0. **(B)** Propidium iodide diffusion assay in lines of *5x per* expressing *pPER64::PER64-mCherry-4G.* Two independent transformant lines, #5-1 and #14-8, are compared to WT and *5x per-2* plants. Numbers indicate the endodermal cell in which apoplastic block is established, counting from the onset of endodermis cell elongation (mean, n=10 seedlings per condition; one-way ANOVA, and Tukey’s test as post hoc analysis; ****P<0.0001; **P<0.005; NS, not significant).

**Supplementary figure 11.**
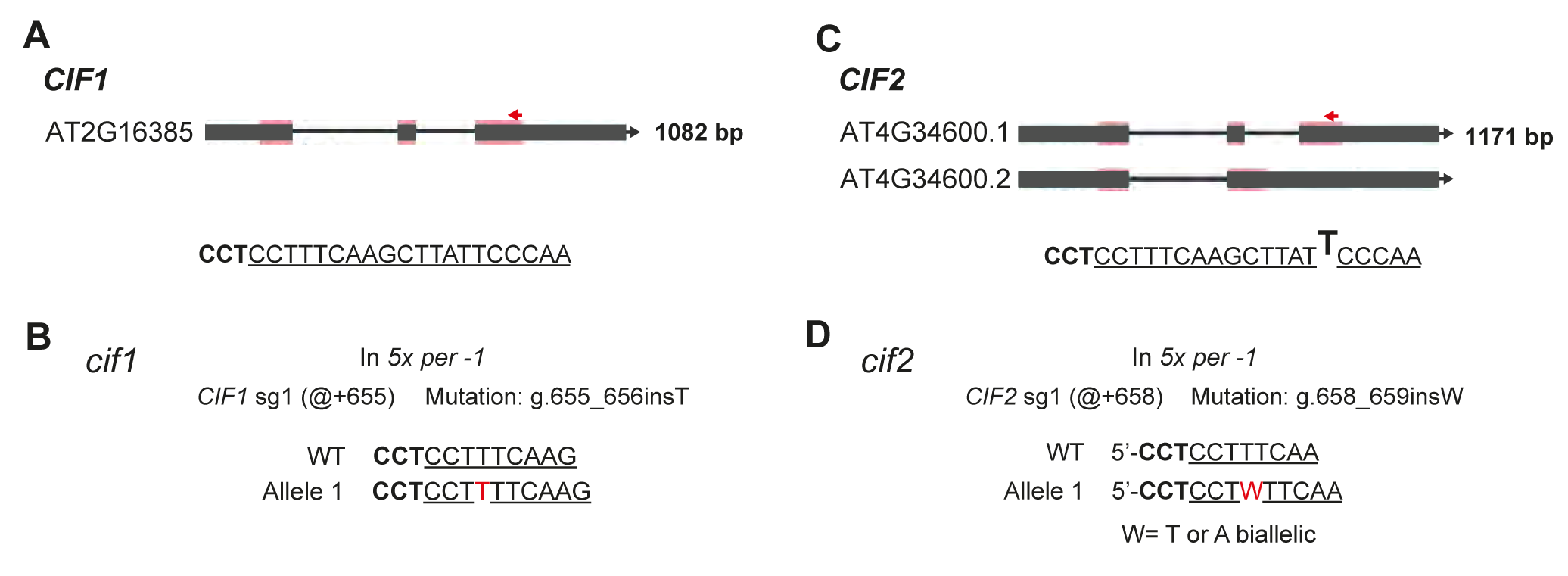
Generation of *cif1* and *cif2* double mutant in *5x per −1* plants by CRISPR-Cas9 gene editing. **(A** and **C)** Schematic of the *CIF1* and *CIF2* genes and CRISPR spacer. The spacer anneals with one mismatch to the target site in *CIF2*. This is represented by “T” in bold text in the sequence of the spacer. Dark grey rectangle, transcribed region; dark-grey rectangle with pink layers, coding DNA sequences; lines, introns; red arrow, localization of target loci; underlined sequence, CRISPR spacer region; bold text, PAM sequence. **(B)** One *cif1* mutation was obtained in the quintuple mutant *per3;9;39;64;72* (*5x per −1*). @+ symbol on wild-type Col-0 (WT) sequence, position of mutation counting from the start codon. Red characters in allele sequences indicate gene mutations. Description of mutation following the nomenclature in (6). **(D)** One biallelic mutant of *cif2* was obtained. Symbols same as in **B**.

**Supplementary figure 12.**
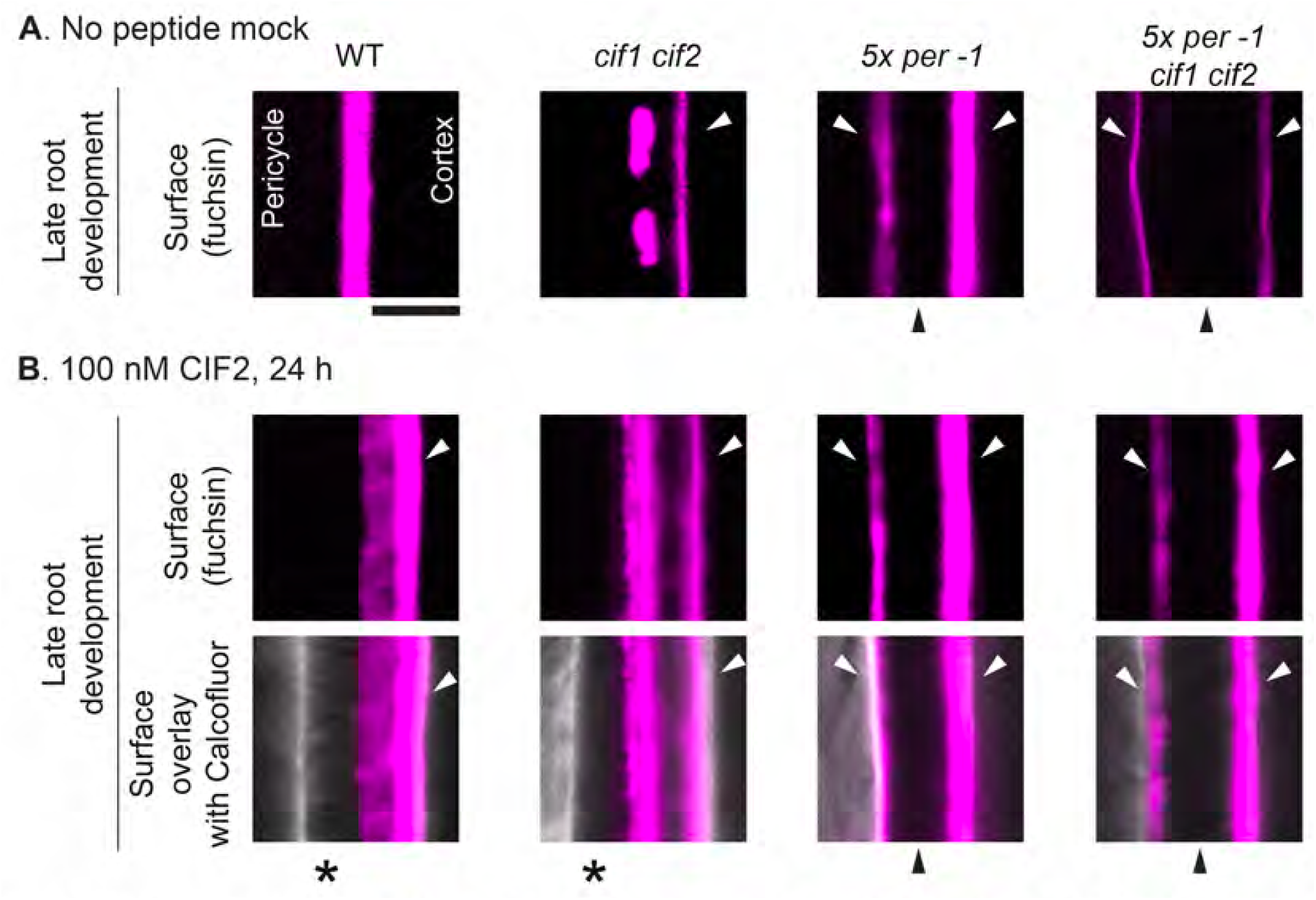
Ectopic deposition of lignin-like material in *5x per* (*3;9;39;64;72*) plants is dependent on CIF1 and 2 small peptides. **(A)** Surface confocal sections of the Casparian strip (CS) stained with basic fuchsin. Pictures were taken at late root development (at metaxylem lignification). WT, wild-type Col-0. **(B)** Same type of pictures were taken in roots of seedlings treated with 100 nM CIF2 for 24 h. Black arrowheads point to the cell wall position where the CS usually forms. White arrows point to ectopic basic fuchsin-positive deposits (lignin-like material). Asterisks indicate the pericycle-facing domain of the cell wall in the endodermis. Fluorescent lignin was stained with basic fuchsin and cell wall was counterstained with calcofluor white M2R following a modified version of the Clearsee protocol (7,8). Scale bar, 5 μm.

**Table S1.**
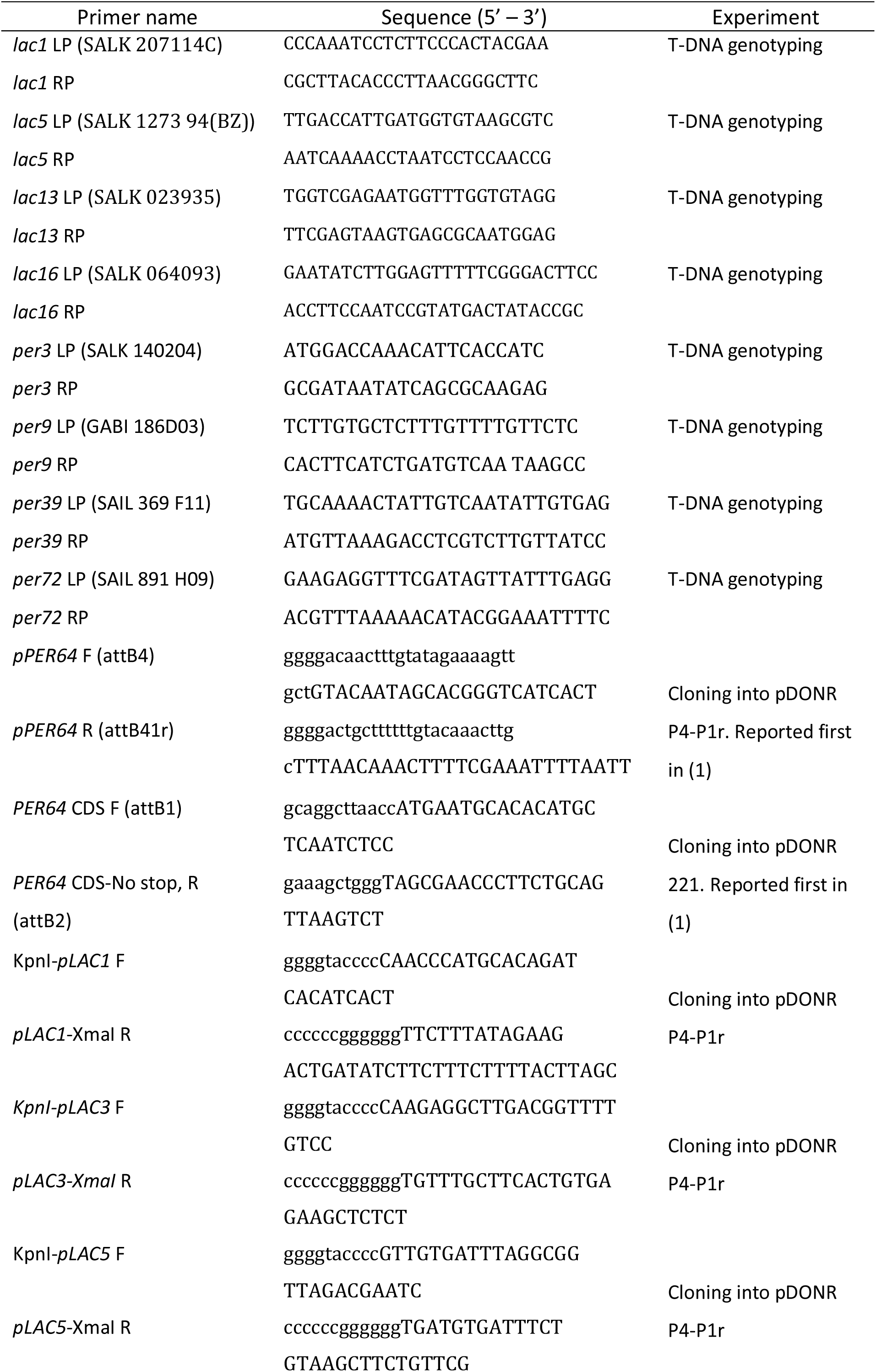
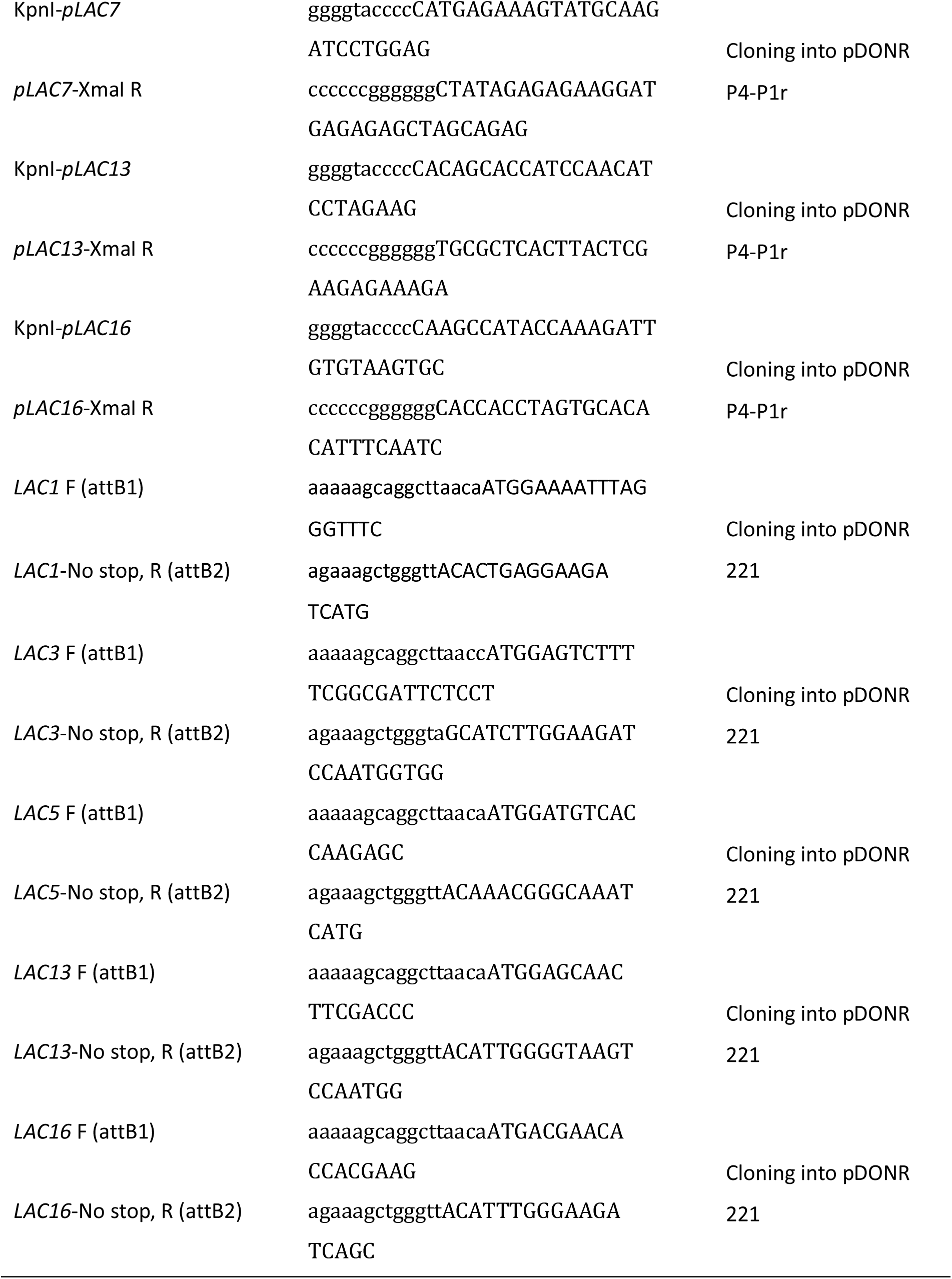
Oligonucleotides used in this study.

**Table S2.**
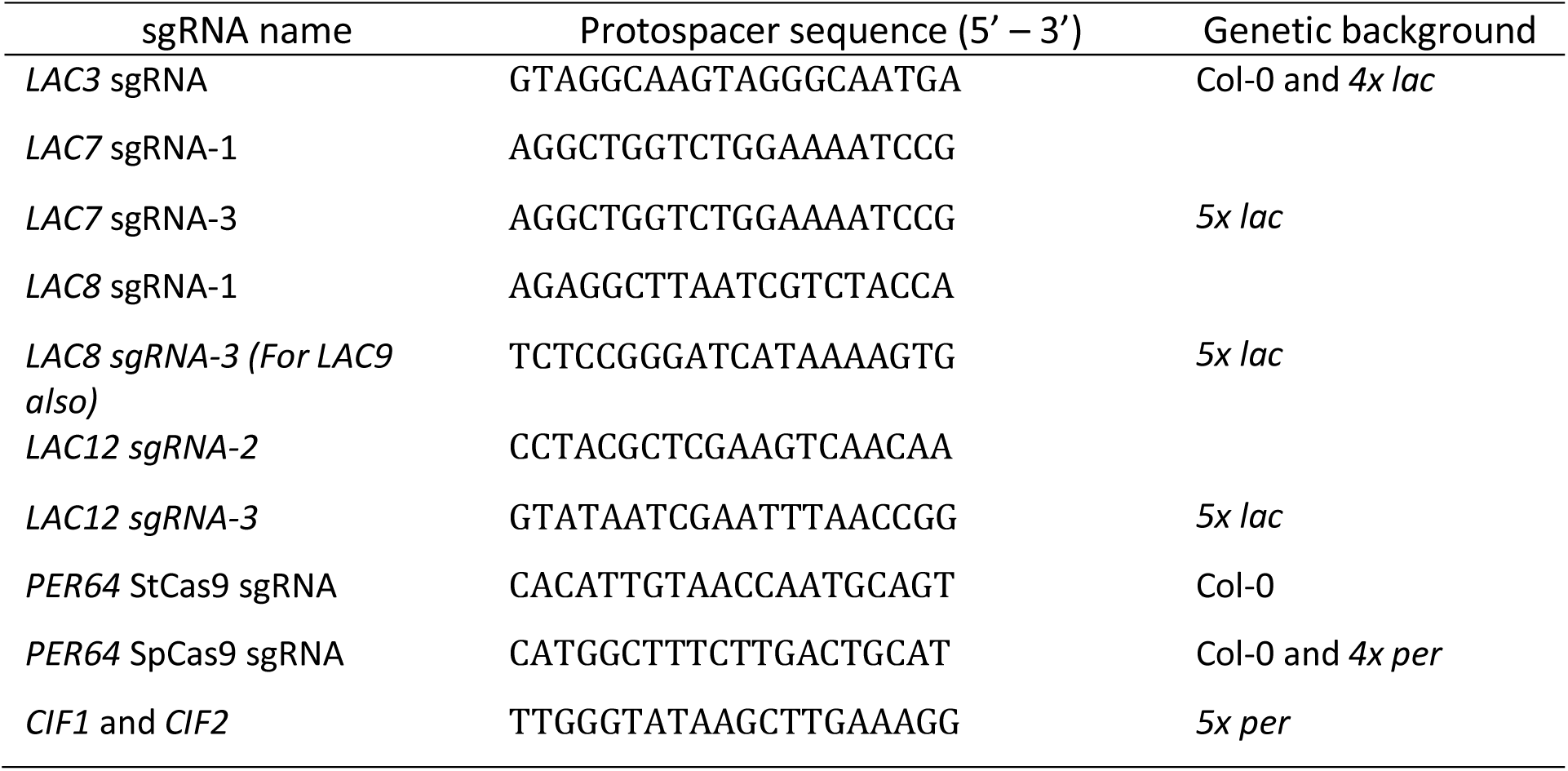
sgRNA protospacers for CRISPR-Cas9 used in this study.

